# Single-cell transcriptome maps of myeloid blood cell lineages in *Drosophila*

**DOI:** 10.1101/2020.01.15.908350

**Authors:** Bumsik Cho, Sang-Ho Yoon, Daewon Lee, Ferdinand Koranteng, Sudhir Gopal Tattikota, Nuri Cha, Mingyu Shin, Hobin Do, Yanhui Hu, Sue Young Oh, Seok Jun Moon, Norbert Perrimon, Jin-Wu Nam, Jiwon Shim

## Abstract

*Drosophila* lymph gland, the larval hematopoietic organ comprised of prohemocytes and hemocytes, has been a valuable model for understanding mechanisms underlying hematopoiesis and immunity. Three types of mature hemocytes have been characterized in the lymph gland: plasmatocytes, lamellocytes, and crystal cells, which are analogous to vertebrate myeloid cells. Here, we used single-cell RNA sequencing to comprehensively analyze heterogeneity of developing hemocytes in the lymph gland, and discovered novel hemocyte types, stem-like prohemocytes, and intermediate prohemocytes. Additionally, we identified the emergence of the lamellocyte lineage following active cellular immunity caused by wasp infestation. We unraveled similarities and differences between embryonically derived- and larval lymph gland hemocytes. Finally, the comparison of *Drosophila* lymph gland hemocytes and human immune cells highlights similarities between prohemocytes and hematopoietic stem cell, and between mature hemocytes and myeloid cells across species. Altogether, our study provides detailed insights on the development and evolution of hematopoiesis at single-cell resolution.

## INTRODUCTION

Blood cells are highly specialized cells that play crucial roles such as elimination of foreign threats during immune responses and retaining memories of immunological events (Sakaguchi et al., 2010). In vertebrates, the multifaceted immune system is composed of two lineages, phagocytic myeloid and memory-dependent lymphoid lineages, to allow holistic and cooperative defense of animals (Weissman et al., 2001). Blood cells in *Drosophila*, collectively called hemocytes, are reminiscent of myeloid-lineage blood cells in vertebrates (Banerjee et al., 2019; Crozatier and Vincent, 2011; Gold and Bruckner, 2014), and are represented by at least three morphologically distinct hemocyte populations: plasmatocytes (PM), crystal cells (CC), and lamellocytes (LM).

Plasmatocytes, which comprise ∼95 % of the hemocytes, play a role in phagocytosis, tissue remodeling, and cellular immune responses – much like macrophages, their vertebrate counterparts (Franc et al., 1996; Irving et al., 2005; Kocks et al., 2005; Kurucz et al., 2007). Crystal cells account for ∼5 % of the blood population and are characterized by crystalline inclusions made up of prophenoloxidase (ProPO), an enzyme responsible for activating the melanization cascade (Binggeli et al., 2014; Gajewski et al., 2007; Lebestky et al., 2000; Tang et al., 2006). Finally, lamellocytes, which are seldom seen in healthy animals grown at normal conditions, mostly differentiate upon parasitic wasp infestation or environmental challenges (Anderl et al., 2016; Honti et al., 2014; Rizki and Rizki, 1984; Sorrentino et al., 2007; Xavier and Williams, 2011).

Blood development in vertebrates involves the primitive and definitive waves of hematopoiesis (Galloway and Zon, 2003). Reminiscent of vertebrate hematopoiesis, two hematopoietic waves have been described during *Drosophila* development, embryonic and larval definitive hematopoiesis (Evans et al., 2003; Hartenstein, 2006). Embryonic hematopoiesis initiates in the head mesoderm of stage-7-embryos and gives rise to hemocytes that migrate throughout the embryo (Holz et al., 2003; Tepass et al., 1994). Upon hatching, embryonically derived hemocytes spread throughout the larva with one hemocyte population freely circulating in the hemolymph and the other colonizing local microenvironments including segmentally repeated hematopoietic pockets of the larval body wall (Leitao and Sucena, 2015; Makhijani et al., 2011).

Definitive hematopoiesis is initiated from hemangioblast-like cells in the embryonic cardiogenic mesoderm, which give rise to the larval lymph gland (Mandal et al., 2004). Medially located prohemocytes, which sustain the developmental potential to give rise to all three mature hemocyte types, constititute the medullary zone (MZ) and continue to proliferate until the early third instar (Jung et al., 2005). Mature hemocytes emerge at the distal edge of the lymph gland, the cortical zone (CZ), from mid-second instar (Kocks et al., 2005; Kroeger et al., 2012). Located between the undifferentiated MZ and the differentiated CZ, is the intermediate zone (IZ) that contains a group of differentiating cells expressing markers for both the MZ and the CZ (Krzemien et al., 2010). The posterior signaling center (PSC), a small group of cells that secrete various ligands, regulates proper growth of the rest of the lymph gland (Benmimoun et al., 2015; Krzemien et al., 2007; Mandal et al., 2007). Lymph glands from healthy larvae reared under normal lab conditions generally follow fixed developmental states until late third instar. Remarkably, following the onset of pupariation, the lymph gland disintegrates, allowing hemocytes to disperse into circulation (Grigorian et al., 2011; Regan et al., 2013).

Female wasps attack second-instar larvae via a sharp needle-like ovipositor that efficiently delivers their eggs (Lemaitre and Hoffmann, 2007; Yang et al., 2015). Wasp eggs trigger cellular immune responses that accompany lamellocyte differentiation in both embryonic and lymph gland hemocytes. Lamellocytes are seen in circulation by 24 hours post-infestation, whereas lymph gland hemocytes remain intact in their original location. Within 48 hours after infestation, a massive differentiation of lamellocytes takes place followed by disruption of the lymph gland (Markus et al., 2009; Sorrentino et al., 2002; Tokusumi et al., 2009). Hemocytes in the lymph gland eventually dissociate into circulation, and mature lamellocytes derived from the lymph gland and hematopoietic pockets encapsulate and neutralize wasp eggs (Irving et al., 2005; Lanot et al., 2001).

The *Drosophila* lymph gland has been largely characterized based on genetic markers and cellular morphology. However, the molecular underpinnings of hematopoietic cells such as different states and the gene regulatory network of each cell type have been less investigated. In addition, questions as to how prohemocytes and mature hemocytes differentiate into lamellocytes upon active immunity, and to what extent hemocytes derived from the embryonic and the lymph gland hematopoiesis differ have been unanswered. Furthermore, due to the lack of sufficient molecular and genetic information, the similarity between *Drosophila* hemocytes and vertebrate immune cells remains to be clarified.

Here, we build an atlas of myeloid-like *Drosophila* hemocytes by taking advantage of single-cell RNA sequencing (scRNA-seq) technology and establish a detailed map for larval hemocytes in the developing lymph gland. We uncover novel classes of hemocytes and their differentiation trajectories, and describe molecular and cellular changes of myeloid hemocytes upon immune challenges. Furthermore, we identify both distinct and common characteristics of hemocytes originating from embryonic and definitive lineages. We also document the evolutionary similarities between *Drosophila* hemocytes and human immune cells. Altogether, our study will stimulate future studies on the function and evolution of the myeloid blood cell lineage.

## RESULTS

### Single-cell transcriptomic profiling of developing myeloid hemocytes

The lymph gland is the larval hematopoietic organ composed of multiple hemocyte cell types and states (Banerjee et al., 2019; Crozatier and Vincent, 2011). To understand the cellular diversity of developing myeloid-like hemocytes in *Drosophila* lymph glands at a single-cell level, we dissected and dissociated lymph glands at three developmental time points, 72, 96, and 120 hours after egg laying (AEL), and applied single cells to Drop-seq, a droplet-based microfluidics platform (Macosko et al., 2015) (Figure 1A). Fourteen independent sequencing libraries, representing 5 each for 72 and 96 h AEL and 4 for 120 h AEL, were prepared for scRNA-seq. We integrated the sequencing libraries after correcting for batch effects within and between time points using Seurat3 (Butler et al., 2018; Stuart et al., 2019). Our quality-control pipeline eliminated cells with outlier unique molecular barcode (UMI) counts, low gene numbers, high mitochondrial gene contents as well as doublets predicted by Scrublet (see Methods for details). As a result, a total of 22,645 cells (72 h AEL: 2,321; 96 h AEL: 9,400; 120 h AEL: 10,924 cells) were retained for subsequent analyses (Figure S1A). The number of cells yielded 5.5 X, 6.8 X, and 2.4 X cell coverage of one lymph gland lobe at 72, 96, and 120 h AEL, respectively (Figure 1B). The 22,645 high-quality cells exhibited a median of 6,361 transcripts (UMIs) and 1,477 genes per cell (Table S1; Figure S1B-S1C). In addition, the scRNA-seq libraries of individual time points corresponded well with genes detected in bulk RNA-seq (≥ 1 TPM; 8,724, 7,654, and 7,627 genes at 72, 96, and 120 h AEL, respectively), while undetected remainders displayed low expression levels (Figure S1D). Furthermore, we validated that the scRNA-seq libraries from 72, 96, or 120 h AEL align well with the pseudotime array of each library (Figure S1E). Altogether, our libraries appear sufficiently complex to reflect the whole transcriptome of developing hemocytes including minor cell types.

**Figure 1.**
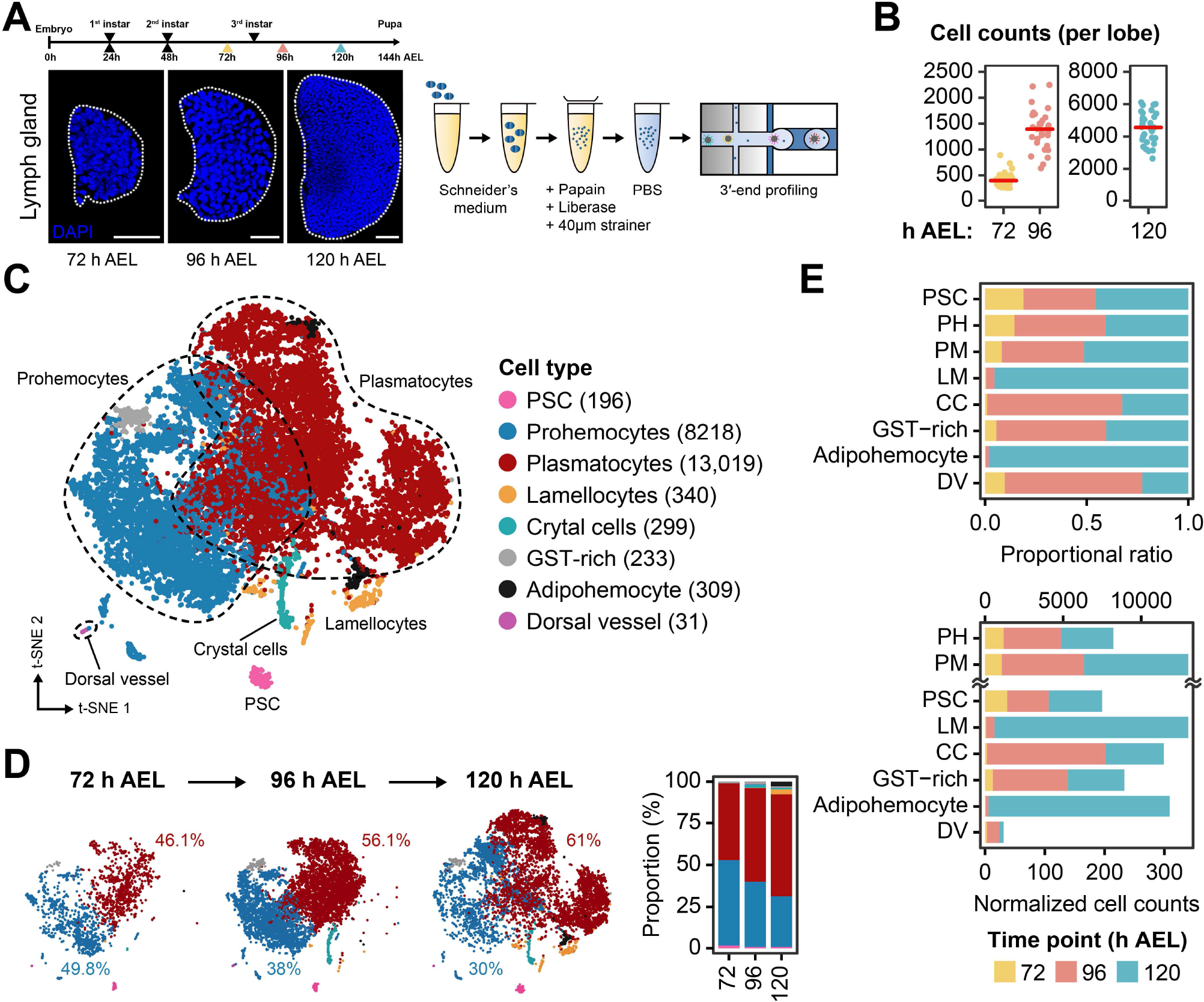
Major cell types identified in developing *Drosophila* lymph glands. (A) *Drosophila* lymph glands (blue, DAPI) at three timepoints (72, 96, and 120 h AEL; After Egg Laying) (left). Schematic workflow of sample preparation for scRNA-seq using Drop-seq (right). Scale bar, 30 μm. Lymph glands are demarcated by white dotted lines. (B) DAPI-positive cell counts of a single lymph gland lobe (*n* = 30 each for three time points). Red horizontal lines show median counts (397, 1392, and 4557 for 72, 96, and 120 h AEL, respectively). (C) A *t*-SNE plot showing the two-dimensional projection of eight major cell types identified in the scRNA-seq dataset (*n* = 22,645). The count of each cell type is indicated in parentheses. Colors denote cell types. Dotted lines demarcate prohemocytes (blue) and plasmatocytes (red). (D) Two-dimensional projections of major cell types along developmental time points (left) and proportion of the cell types at each time point (right). Proportions of prohemocytes (blue) and plasmatocytes (red) are indicated. (E) Relative proportion (indicated as proportional ratio; top) and normalized cell counts (bottom) of each major cell type. Colors represent sampling time points.

### Major cell types and transcriptional dynamics of *Drosophila* hematopoiesis

After validating the quality of the single cell libraries, we mapped the cells to the major zones of the lymph gland (Jung et al., 2005). To explore the major cell types in the developing lymph gland, we aligned cell clusters from the three developmental time points using the Louvain algorithm (Blondel et al., 2008) and visualized the data using a nonlinear dimensionality reduction by *t*-distributed stochastic neighbor embedding (*t-*SNE) (van der Maaten and Hinton, 2008). We aggregated cell clusters according to the expression of previously published marker genes by manual curation and identified seven distinctive groups of isolated hemocytes (Figure 1C; Figure S1F-1G; Table S2). These clusters include prohemocytes (PH: *Tep4*, *Ance*; 36.2 %), plasmatocytes (PM: *Hml*, *Pxn*, *NimC1*; 57.6 %), crystal cells (CC: *lz*, *PPO1*, *PPO2*; 1.3 %), lamellocytes (LM: *atilla*; 1.5 %), the PSC (*Antp*, *col*; 0.9 %), and two additional clusters with uncharacterized genetic features. One novel cluster is enriched with glutathione-S-transferase transcripts including *GstD1*, *GstD3*, *GstE1, GstE7,* and *GstT1*, that we named “GST-rich” (1.0 %). The other novel cluster exhibits high expression levels of phagocytosis receptor-, lipid metabolism-related, and starvation-induced genes such as *crq, eater*, *Sirup*, *LpR2*, and *Lsd-2*. We referred to this cluster as “adipohemocyte” (1.4 %) based on the name of similar hemocytes in other insects (Hillyer et al., 2003). We verified the presence of GST-rich and adipohemocyte cell populations in wild-type lymph glands by confirming the expression of signature genes for these clusters in matched bulk RNA-seq (Figure S1H). Additionally, we identified cells of the dorsal vessel (DV; 0.1 %) as an extra non-hemocyte cell type based on previously identified marker genes, *Mlc1* and *Hand*, for this tissue (Figure 1C; Figure S1G).

Separation of clustered cells by developmental time points revealed that the relative population size of cell clusters changes as the lymph gland matures. Hemocytes in the lymph gland at 72 h AEL are subdivided into two major groups—prohemocytes and plasmatocytes, with a virtually identical ratio of 49.8 % and 46.1 %, respectively (Figure 1D-1E). As the lymph gland matures, the proportion of plasmatocytes exceeds that of prohemocytes, and only 30 % of the hemocytes retain the prohemocyte signature at 120 h AEL (Figure 1D-1E). This result is consistent with proportional changes of prohemocytes or plasmatocytes visualized by marker gene expression during development *in vivo* (Figure S1F). Different from plasmatocytes emerging from 72 h AEL, crystal cells and GST-rich cells first appear at 96 h AEL, and lamellocytes and adipohemocytes appear later at 120 h AEL (Figure 1D-1E). The PSC maintains constant cell numbers and relative ratios across lymph gland development (Figure 1E). Due to temporal discrepancies in the emergence of mature hemocytes, we observed that most cells at 72 h AEL overlap well with cells in subsequent time points, while cells at 96 and 120 h AEL segregate from those of preceding points on the *t*-SNE plot (Figure S1I). These results were reproduced by an independent analysis using UMAP and an alternative batch correction method (Figure S1J, see Methods for more details).

To better characterize the major cell types and transitions in gene regulatory networks during lymph gland development, we applied SCENIC, an algorithm that reconstructs gene regulatory networks and identifies cell states from scRNA-seq data (Aibar et al., 2017). We identified previously recognized transcriptional regulators such as *jumu*, *Antp*, and *kn* (also known as *collier*) in the PSC; *srp* in plasmatocytes; and *lz* and *Su(H)* in crystal cells (Figure S1K) (Crozatier et al., 2004; Hao and Jin, 2017; Lebestky et al., 2003; Mandal et al., 2007). Moreover, we characterized transcription factors in well-known complexes or pathways in each cell type. In prohemocytes, we detected transcription factors of DREAM (Sadasivam and DeCaprio, 2013), a protein complex responsible for cell cycle regulation, including *E2F2* and *Dp,* and the Dpp pathway transcription factor *Mad* at 96 to 120 h AEL (Figure S1K). Plasmatocytes, on the other hand, utilize distinctive transcriptional regulators of the ecdysone pathway highlighted by *br*, *EcR, usp, Eip74EF,* and *Hr4,* and stress responsive genes such as *foxo* and *dl* (Figure S1K). Overall, our single-cell dataset of the entire lymph gland reliably reveals seven major types of hemocytes (PSC, prohemocytes, plasmatocytes, crystal cells, lamellocytes, and two newly identified populations—GST-rich and adipohemocytes). Also, SCENIC analysis delineates transcriptional transitions of the hemocytes and their regulators at the single-cell level.

### Heterogeneous populations of lymph gland hemocytes

Our scRNA-seq data prompted us to further catalog the heterogeneity of primary cell types by performing unsupervised clustering. With subclustering analysis, we identified eleven subclusters of prohemocytes, ten subclusters of plasmatocytes, and two subclusters each for lamellocytes and crystal cells (Figure 2A; Figure S2A). We ensured that each subcluster contains cells from all libraries except stage-specific subsets (Figure S2B). In addition, we excluded non-hemocyte subclusters enriched with ring gland- or neuron-specific genes (Figure S2C-S2D). Adipohemocytes split into three subordinate clusters; yet, two subclusters were library specific, and thus, only one subcluster was considered relevant and kept for subsequent analyses (Figure S2B). Interestingly, both the PSC and GST-rich clusters did not split into subclusters (Figure S2A).

**Figure 2.**
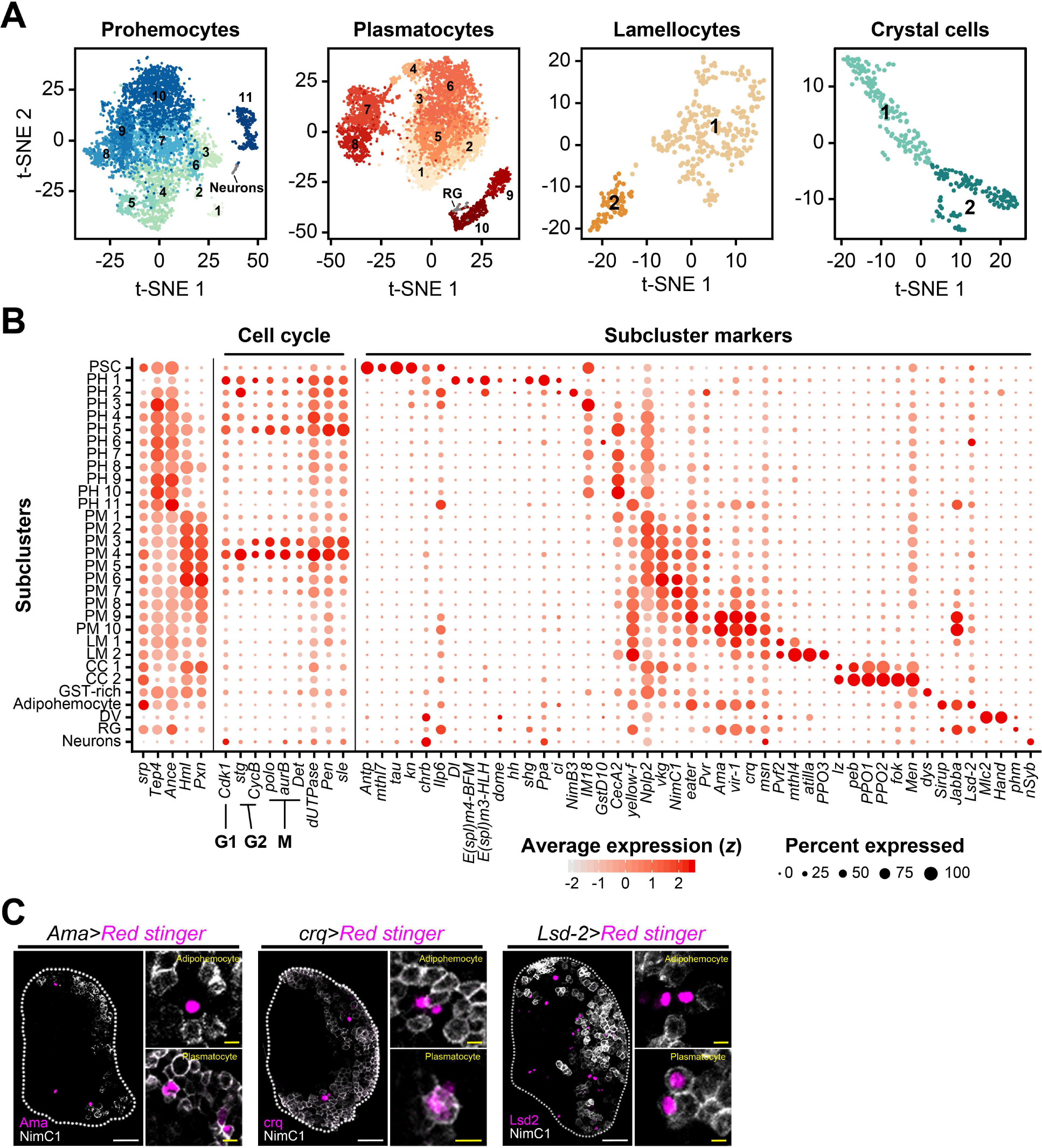
Heterogeneous cellular states of hemocytes in the lymph gland defined by subclustering analysis. (A) Subclusters of hemocytes—prohemocytes, plasmatocytes, lamellocyte, and crystal cells—are projected onto two-dimensional *t*-SNE plots. Non-hematopoietic cell types (Neurons; RG, ring gland) are indicated. The numbers in the plots represent the subcluster number. (B) Dot plot presentation of significant gene sets in the 31 subclusters. 5 representative markers, *srp*, *Tep4*, *Ance*, *Hml*, and *Pxn* are indicated to the left column. Cell-cycle regulating genes are shown in the middle. Signature genes identified in this study are marked with subcluster markers. Dot color shows levels of average expression, and dot size represents the percentage of cells expressing the corresponding marker genes in each subcluster. (C) *Ama* (magenta; *Ama-Gal4 UAS-Red stinger*), *crq* (magenta; *crq-Gal4 UAS-Red stinger*) or *Lsd-2* (magenta; *Lsd2-Gal4 UAS-Red stinger*) partially overlap with NimC1 (white). *Ama^+^NimC1^+^, crq^+^NimC1^+^* and *Lsd-2^+^NimC1^+^* cells correspond to plasmatocytes, whereas *Ama^+^NimC1^-^, crq^+^NimC1^-^* and *Lsd-2^+^NimC1^-^* cells represent putative adipohemocytes. Generally, *Ama^+^NimC1^-^, crq^+^NimC1^-^* and *Lsd-2^+^NimC1^-^* cells show higher *Ama*, *crq*, or *Lsd-2* expression levels than those of double-positive cells. Magnified images show colocalization of *Ama*, *crq or Lsd2* with NimC1 (right in each panel). White scale bar indicates 30 μm; yellow scale bar, 3 μm. White dotted line demarcates the lymph gland.

The majority of prohemocyte subclusters is evenly represented at all timepoints and maintains high levels of *Tep4* and *Ance* throughout (Figure 2B). Apart from their constant expression, we identified discrete fluctuations of cell-cycle regulating genes including *polo*, *Cdk1*, *aurB*, *Det*, *CycB*, and *stg* within prohemocytes, accompanied by alterations in additional nuclear genes such as *dUTPase*, *Pen*, and *sle* (Figure 2B). These genes peak at PH1, PH2, and PH4–PH5 within prohemocytes, and a comparable pattern is also present in PM3–PM4 (Figure 2B). There are obvious distinctions between genes involved in the cell cycle: *stg* and *CycB* are regulators of the G2 phase; *Cdk1* of the G1 phase; and *polo*, *aurB*, and *Det* are regulators of the M phase (Edgar et al., 1994; Llamazares et al., 1991; Mathieu et al., 2013; Parry and O’Farrell, 2001). Based on relative levels of these genes, PH1 is likely to be in G2 and M; PH2 in G2; PH4 in G1; PH5 in M; and PM3 and PM4 in M phase (Figure 2B). Similar to prohemocytes, plasmatocytes exhibit gradual changes in *vkg*, *NimC1,* and *eater* while keeping *Hml* and *Pxn* expression high (Figure 2B). PM7 to PM10 express characteristic signature genes such as *Ama*, *vir-1*, and *crq*, only detected at 120 h AEL (Figure 2A, Figure S2B). Crystal cells are divided into two groups, CC1 and CC2 (Figure 2A). CC1 expresses low levels of *lz* along with the expression of MZ and CZ markers. However, CC2 is devoid of the MZ or CZ markers and only displays high levels of *PPO1* and *PPO2*, suggesting that CC1 and CC2 correspond to early and mature crystal cells, respectively (Figure 2A-2B). Similarly, lamellocytes are separated into premature LM1 and mature LM2 reminiscent of the CC1 and CC2 clusters (Figure 2A-2B).

We next sought to identify new markers and characteristic gene expression patterns in the lymph gland. We confirmed the expression of *Ilp6*, *tau*, *mthl7*, *brat,* and *chrb* in the PSC; *Men* and *Numb* in crystal cells; and *vir-1* and *Ama* in plasmatocytes (Figure 2B, Table S2; Figure S2E-S2G). In addition, we discovered that *zfh1* and *tep2* are expressed in prohemocytes in addition to representative markers such as *Ance* and *dome* in the MZ (Figure S2H-S2I). *crq*, *vir-1*, and *Ama* are significantly expressed in both adipohemocytes and mature plasmatocytes; however, adipohemocytes exhibit high levels of *crq* and *Lsd-2* while keeping low levels of *NimC1* (Figure 2C). In addition to markers widely used (Evans et al., 2014; Yu et al., 2018), we identified new enhancer-trap or MiMIC lines (Nagarkar-Jaiswal et al., 2015) targeting the lymph gland marker genes (Table S3). Lastly, we confirmed the mRNA expression pattern of previously reported genes in each subset (Banerjee et al., 2019) (Figure S3J). Together, we classified 28 transcriptionally distinctive subtypes of hemocytes in the developing lymph gland and assigned functional descriptions of each subset based on gene expression patterns. *Bona fide* markers elucidated in each subcluster collectively provide a valuable resource for further understanding of myeloid hemocyte development.

### Trajectory reconstitution and functional networks

Hematopoietic events involving transitions of hemocytes from their stem-like to final cell types have been a major focus for understanding *Drosophila* hematopoiesis. Thus, we investigated the time sequence of lymph gland hematopoiesis by reconstruction of developmental trajectories using Monocle 3 (Cao et al., 2019). For the trajectory analysis, we excluded PSC as the PSC cells do not give rise to the rest of lymph gland hemocytes (Figure S3A) (Crozatier et al., 2004; Mandal et al., 2007), and we set the PH1 subcluster as the start point based on the expression of *Notch*, *shg*, and high levels of mitotic genes (Figure 2B). Pseudotime reconstitution of lymph gland hemocytes displays the main trajectory from prohemocytes to plasmatocytes along with divergent sub-trajectories towards crystal cells, adipohemocytes, GST-rich, and lamellocytes (Figure 3A-3B; Video S1). The trajectory corresponds well with on-and-off patterns of marker genes in the lymph gland (Figure S3B). Moreover, there is an excellent correlation between the real-time and the pseudotime trajectories when compared with segregated real-time hemocyte transcriptomes (Figure 3B-3C, Figure S5C). These analyses validate the *in silico* algorithm-based sequence of hemocyte differentiation and confidently illustrate the developmental phases of lymph gland hemocytes.

**Figure 3.**
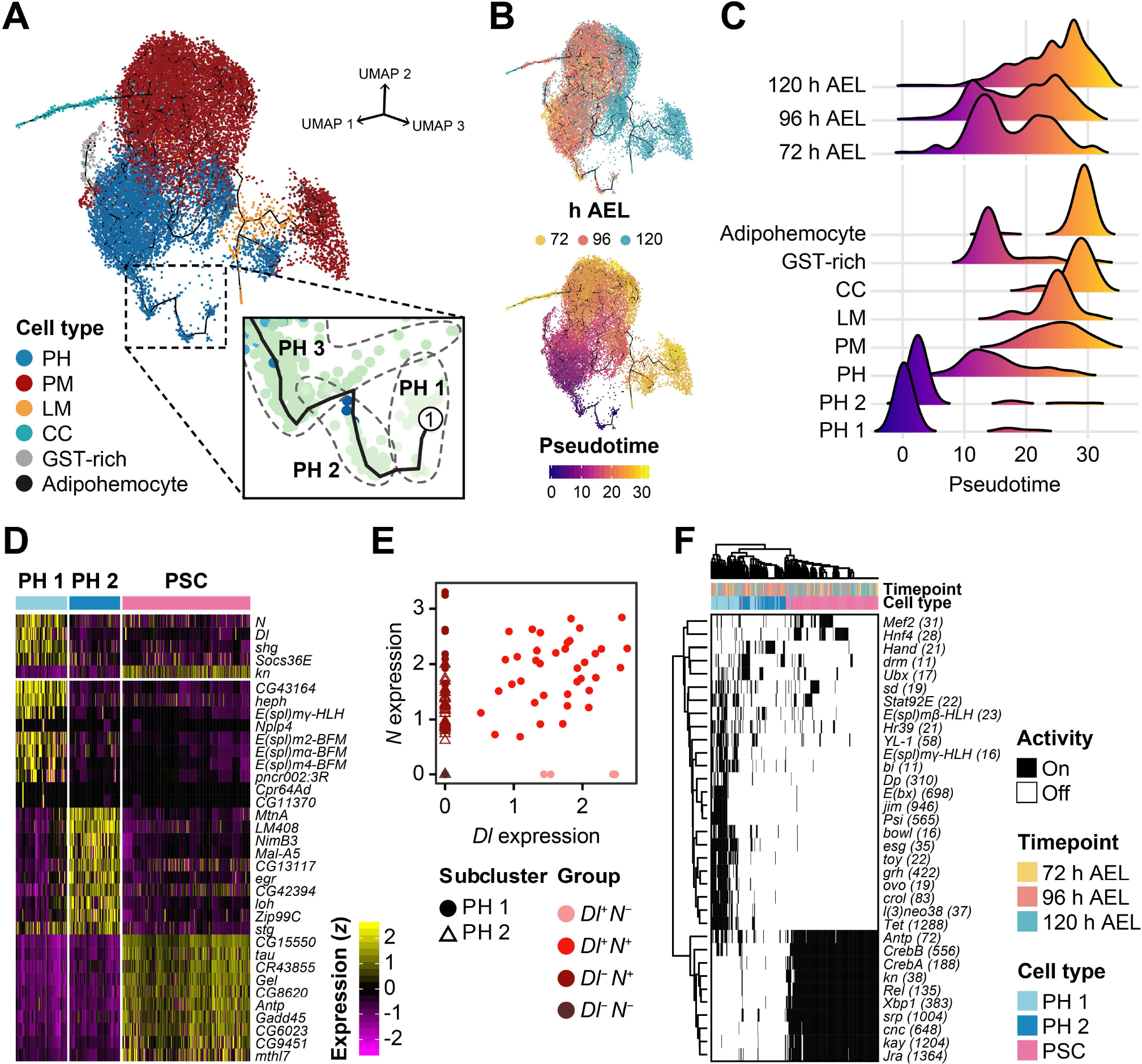
Reconstruction of lymph gland hematopoiesis using pseudotime trajectory analysis. (A) A three-dimensional landscape of the lymph gland hematopoiesis trajectory using Monocle 3 (*n* = 19,143). Non-hematopoietic cells were excluded in this analysis. Black line indicates the trajectory. Colors indicate the six major cell types used for analysis. The inset shows the three ancestral PH subclusters, PH1, PH2, and PH3. (B) Trajectories re-drawn by developmental time points (top) and calculated pseudotime (bottom). Colors indicate the three real-time points (top) and pseudotime (bottom). (C) Relative densities of hemocytes segregated by three time points (top) and cell types (bottom) along pseudotime. PH1 and PH2 are separated from other PH subclusters for higher resolution. Colors in density plots correspond to pseudotime, as in B. (D) Heatmap representation of the 35 signature genes identified in PH1, PH2, and the PSC (*n* = 77, 79, and 189 cells, respectively). The colored legend denotes the standardized level of the genes. (E) Four subgroups in PH1 and PH2 defined by the expression of *Delta* (*Dl*) and *Notch* (*N*). Colors show subgroups and shapes specify PH subclusters. X axis means *N* expression; Y axis, *Dl* expression. (F) Binary heatmap showing the activity of transcription factors in PH1, PH2, and the PSC predicted by SCENIC. Numbers in parentheses denote the count of downstream genes used to test the activity of transcription factors.

In the major trajectory, we observed a linear trajectory from PH1 to PH3, projecting towards diverse differentiating states of prohemocytes including PH4-PH8 and GST-rich (Figure 3A, Figure S3D-S3E). In the later sequence, all the PH subclusters including the GST-rich merge into PM1 in the trajectory, implying that GST-rich is a tributary of prohemocytes joining the main PH-to-PM flow (Figure S3D-S3E). A branch is observed following PM1, producing separable paths towards either the plasmatocyte, the crystal cell, or the lamellocyte lineages. PM3 is biased to the plasmatocyte and the crystal cell lineages, while PM4 gives rise to late plasmatocytes, adipohemocytes, and lamellocytes (Figure S3D-S3E). We also observed a coupling of cell division and differentiation.

Besides PH1 and PH2 subtypes expressing high levels of cell cycle genes, PH4-PH5 and PM3-PM4 emerge immediately after each divergence (Figure 2B; Figure S3D-S3E). As an auxiliary route, PH9 and PH11 are distinguishable from PH3 and bypass the classical PH-to-PM flow to give rise to late plasmatocytes or lamellocytes at 120 h AEL (Figure 3A-3B, Figure S3D-S3E).

To address the functional characteristics of the hematopoietic trajectory and associated subclusters, we examined subtrajectory- and subcluster-specific gene-expression modules to determine whether subclusters share similar gene expression modules (Figure S3F). Strikingly, prohemocyte subclusters exhibit related translation-, metabolism-, and signaling gene expression whereas plasmatocyte subclusters show relatively high levels of extracellular matrix (ECM), cytoskeletal and immune responsive genes. Crystal cell subclusters display high levels of genes involved in sugar metabolism, and adipohemocyte and GST-rich subclusters show fatty acid-related and DNA damage responsive gene modules, respectively (Figure S3F).

We next focused on the transition of prohemocytes into mature hemocytes and defined subclusters spanning the intermediate zone according to the trajectory analysis and modular configurations. Given that PH5 and PM3/PM4 are mitotic and PH6, PH8, PH10, GST-rich, and PM1, subclusters between PH5 to PM3/PM4, exhibit moderate levels of *Tep4*, *Ance*, *Pxn* and *Hml*, we hypothesized that PH6, PH8, PH10, GST-rich, and PM1 correspond to intermediate cell types prior to differentiation. These subclusters are found at all-time points (Figure S2B). We scored highly enriched genes in the potential intermediate subclusters and noticed that expression of *Nplp2* aligns well with these cell populations (Figure 2B). Visualizing *Nplp2* in the lymph gland revealed a partial overlap of *Nplp2* with the MZ marker, *Dome^Meso^*, or the CZ marker, *Pxn*. However, as *Nplp2* is expressed independently from the late plasmatocyte marker, NimC1 (Figure S3G), it indicates that *Nplp2* is expressed in the intermediate zone of the lymph gland, which corresponds to hemocytes in transition towards differentiation. Altogether, the pseudotime trajectory analysis provides a detailed basis for prohemocyte differentiation. In addition, we demonstrate the presence of subclusters in transition, previously described as the IZ, and their endogenous gene expression.

### Initiation of hematopoiesis in the lymph gland

Prohemocytes have been established as the precursors of lymph gland hemocytes that produce the entire lymph gland hemocytes (Jung et al., 2005; Minakhina and Steward, 2010b). Despite previous attempts to understand the onset of larval hematopoiesis, it has been unclear whether there is a premature state of prohemocytes reminiscent of mammalian hematopoietic stem cells (HSCs).

To investigate the primordial cell types during lymph gland hematopoiesis, we focused on the earliest PHs—PH1 and PH2—which initiate the entire trajectory. We observed that the majority of PH1 is detached from PH2/PH3 and a subset is connected to PH2 in the trajectory map (Figure 2A, 3A, Figure S2A, S3B-S3D). Though PH1 and PH2 mark the earliest pseudotime, both clusters are found at all developmental time points (Figure 3B-3C, Figure S2B).

We identified multiple signature genes in the PH1 subcluster (Figure 2B, 3D). First, we discovered that *Notch* (*N*), its ligand, *Delta* (*Dl*), and the *E(Spl)* family genes, downstream targets of the Notch pathway, are expressed in PH1 (Figure 3D, Figure S4A). Interestingly, cells in PH1 and PH2 are sequentially arrayed according to on-and-off patterns of *Dl* and *N* (Figure 3E, Figure S4B). Second, we found associations of Hippo, MAPK, Wnt, and Notch pathways with *Dl^+^N^+^* cells of PH1 by KEGG pathway analysis (Figure S4B). Third, we observed levels of *dome*, *hop*, *Stat92E*, and *Socs36E* in PH1, reflecting active JAK/STAT signaling (Figure 3D, Figure S4A-S4B). Strikingly, the expression of Notch/Delta and JAK/STAT-related genes in PH1 decrease in the succeeding PH2, suggesting that PH1 cells undergo a drastic change. Fourth, we identified that PH1 does not express *col* while PH2 exhibits low levels of *col* expression, consistent with previous observations (Figure S4C)(Benmimoun et al., 2015). Lastly, high levels of cell cycle genes are detected in both PH1 and PH2, constituting some of the few PH subclusters actively proliferating in the lymph gland (Figure 2B).

Next, we applied SCENIC to further establish gene regulatory networks of PH1 and PH2 cell populations with PSC as a comparison. SCENIC analysis on the PSC successfully proved the activity of known transcription factors (Figure 3F). Moreover, the PH1 subcluster displays transcriptional activity of known regulators, such as *sd* and *Stat92E*, as well as novel genes, including *jim*, *Psi*, *bowl*, *esg*, *Tet*, *E(bx)*, and the *E(spl)* family (Figure 3F).

We next performed spatial reconstructions for PH1 *in vivo*, and profiled the expression of genes newly identified in the subset. Interestingly, *Stat92E::edGFP* is expressed in the cells neighboring the PSC, which are neither *Tep4*^+^ MZ nor *Antp*^+^ PSC (Figure 4A, Figure S4D-S4E). Similarly, *Stat92E*^+^ cells show close contact with *col*^+^ cells without having apparent overlaps (Figure 4B). The number of *Stat92E*^+^ cells increases over development, maintaining relatively constant ratios of these cells (Figure 4C). Furthermore, *Stat92E*^+^cells are gone upon genetic ablation of the PSC, which indicates that expression of *Stat92E*^+^ PH1 is dependent upon the PSC (Figure 4D). We additionally detected *Dl* mRNA or Dl protein expression near the PSC similar to the pattern observed with *Stat92E::edGFP* (Figure 4E, Figure S4F). The majority of *Dl^+^* cells are *Stat92E^+^*; however, Dl covers a range broader than a few cell diameters than *Stat92E^+^* (Figure 4F). We screened Gal4 lines of signature genes in PH1 and identified a new *Dl* enhancer-trap that marks *Antp*^-^ cells adjoining to the PSC, which produce hemocytes of the entire lymph gland (Figure 4G). To summarize, PH1, the initial subset of the pseudotime trajectory, indicates a novel subpopulation of prohemocytes that physically interacts with the PSC and is adjacent to *col*^+^ PH2. PH1 cells do not co-localize with conventional MZ or CZ markers but exhibit distinctive gene regulatory networks primarily steered by the Delta/Notch and JAK/STAT pathways. Moreover, these cells retain potentials to give rise to hemocytes in the lymph gland during 72 to 120 h AEL (Figure 4H). Thus, we conclude that PH1 is a premature state of prohemocytes reminiscent of mammalian hematopoietic stem cells (HSCs).

**Figure 4.**
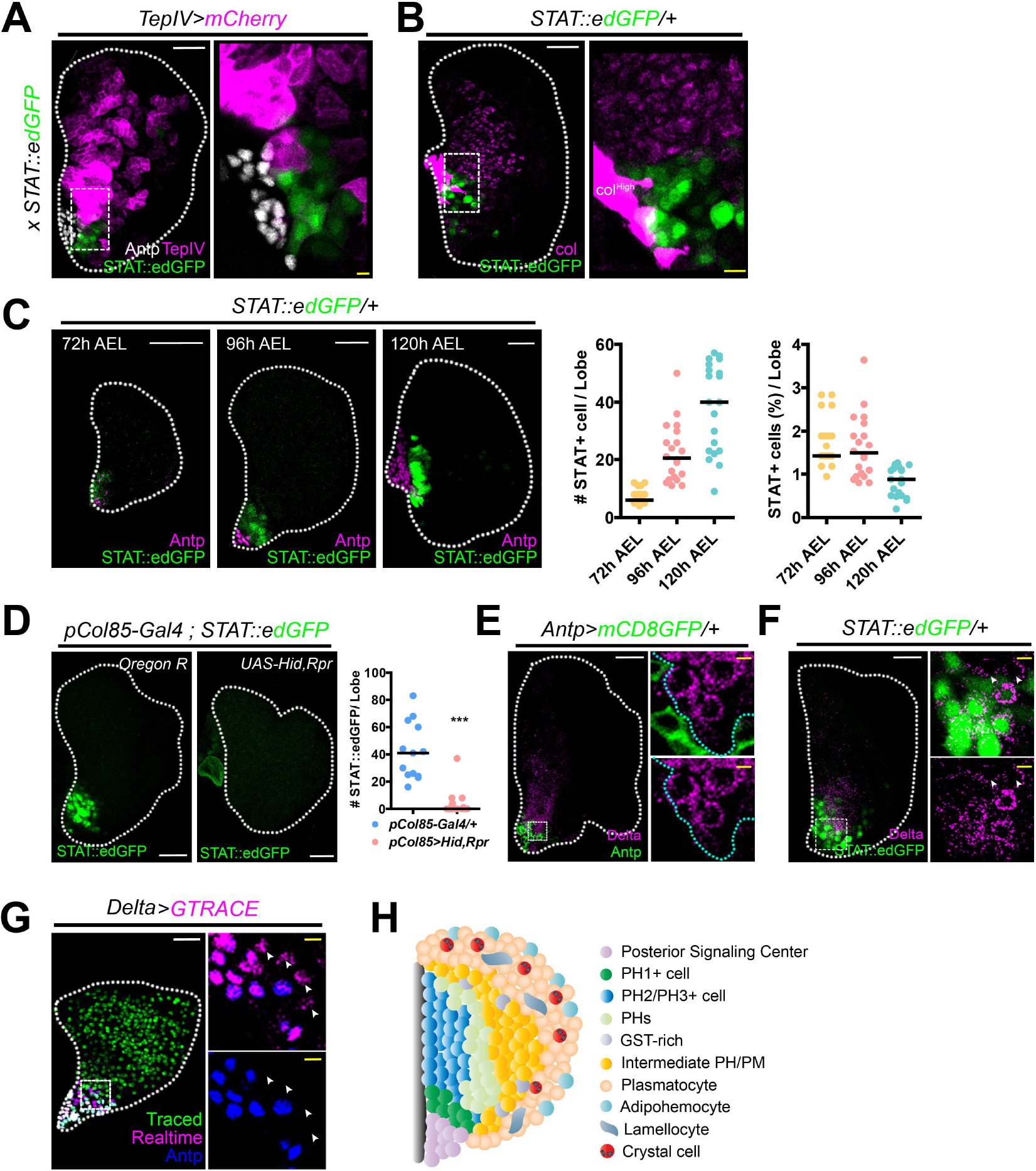
Expression of PH1 in the lymph gland. (A) *STAT92E*^+^ (green) and *TepIV^+^* (magenta) or *Antp^+^* (white) cells are mutually exclusive (*TepIV-Gal4 UAS-mCherry; STAT92E::edGFP*). The dotted box indicates the region magnified. High magnification of *STAT92E*^+^ and *TepIV^+^* cells near *Antp^+^* PSC. (B) *STAT92E^+^* cells (green) do not co-localize with cells expressing high (PSC) or low (PHs) levels of *collier* (magenta). Box indicates magnified view. High magnification of *STAT92E*^+^ and *col^+^* cells near the PSC. (C) The number of *STAT92E*^+^ cells (green) increases during lymph gland development (72, 96, and 120 h AEL). Exclusive expression of *Antp^+^* (magenta) and *STAT92E^+^* (green) is maintained at all time points. Graphs represent quantitation of the number (left) or the proportion (right) of *STAT92E*^+^ cells in one lymph gland lobe. (D) Genetic ablation of the PSC (*pCol85-Gal4; STAT92E::edGFP UAS-hid, rpr*) attenuates *STAT92E* (green) expression in the lymph gland (left). Graph indicates quantitations of the number of *STAT92E*^+^ cells in one lymph gland lobe (right, \*\*\**P*<0.0001). (E) Dl^+^ cells (magenta) are localized adjacent to the PSC (Antp, green). Box indicates the magnified area. Magnified views of Dl^+^ (magenta) and *Antp*^+^ (green) cells (two markers, right top; one marker, right bottom). Cyan dotted lines delineate Dl^+^ expressing cells. (F) Dl^+^ cells (magenta) co-localize with *STAT92E^+^* (green). Box indicates the magnified area. Magnified view of Dl^+^ (magenta) and *STAT92E^+^* (green) cells (right). A few Dl^+^ cells that do not express *STAT92E::edGFP* are indicated (arrowhead). (G) Lineage tracing of *Dl^+^* cells (green, traced; magenta, real time; blue, Antp). *Delta-Gal4 UAS-GTRACE* covers the entire lymph gland. Box indicates the magnified view. Arrowheads represent *Dl^+^* cells next to the *Antp^+^ Dl^+^* PSC. The PSC does not give rise to lymph gland hemocytes (see Supplememtary Figure 3A). (H) Model. PH1 cells are adjacent to the PSC. PH1 and PSC or PH2 are mutually exclusive. There are multiple states of prohemocytes including GST-rich and intermediary PHs/PMs. Plasmatocytes represent an heterogenous cell population including adipohemocytes. Lamellocytes are rarely observed under normal conditions. Crystal cells are found among differentiated plasmatocytes. In panels A through G, white scale bar indicates 30 μm; yellow scale bar, 3 μm. White dotted line demarcates the lymph gland. Median value is represented in graphs (C, D).

### Differentiation of lymph gland hemocytes upon wasp infestation

We next investigated emerging heterogeneity and differentiation of the lymph gland hemocytes upon wasp infestation. We aligned and matched lymph gland hemocytes from 24 h PI (post-infestation; 96 h AEL) to control hemocytes using the label transfer, that resulted in the annotation of six hemocyte clusters (iPSC, iPH, iPM, iCC, iLM, and iGST-rich) and 21 subclusters (9 iPHs, 6 iPMs, 2 iCCs, 2 iLMs, iPSC and iGST-rich) when compared to those from controls (Figure 5A-5B; Figure S5A). Consistent with previous studies (Crozatier et al., 2004; Ferguson and Martinez-Agosto, 2014), wasp infestation significantly reduces crystal cells (iCCs)—iCC1 and iCC2 (Figure 5A-5B, Figure S5B). A similar decline is readily observed in iPH1, which is confirmed by the reduction of *Stat92E*^+^ or *Dl*^+^ iPH1 in the lymph gland upon wasp infestation (Figure 5C, Figure S5B). In contrast, iPH4, iPH6, iPH7, iPM1, and iPM4 show a stark increase in numbers and relative proportions, implying an expansion of differentiating hemocytes upon wasp infestation (Figure S5B). Coinciding with the increase of differentiating cells, the lamellocyte and GST-rich populations, which are barely observed during normal development at 96 h AEL, expand upon wasp infestation (Figure 5A-5B, Figure S5B). Lamellocytes derived upon infestation (iLMs) are subclustered into two groups: iLM1 and iLM2, which represent immature and mature iLMs, respectively (Figure S5C). While other cell types undergo significant modifications upon wasp infestation, we did not detect any changes in the expression and number of iPSC upon wasp infestation (Figure 5B, Figure S5D-S5F). When the signature genes of each subcluster are compared to those from controls, gene expression patterns in general are not altered (Figure S5D). However, the intermediate cell population already expresses lamellocyte markers such as *atilla* and *mthl4*, a novel LM marker (Figure S5D). These data indicate that the active immunity causes a biased commitment of prohemocytes and plasmatocytes to the lamellocyte lineage.

**Figure 5.**
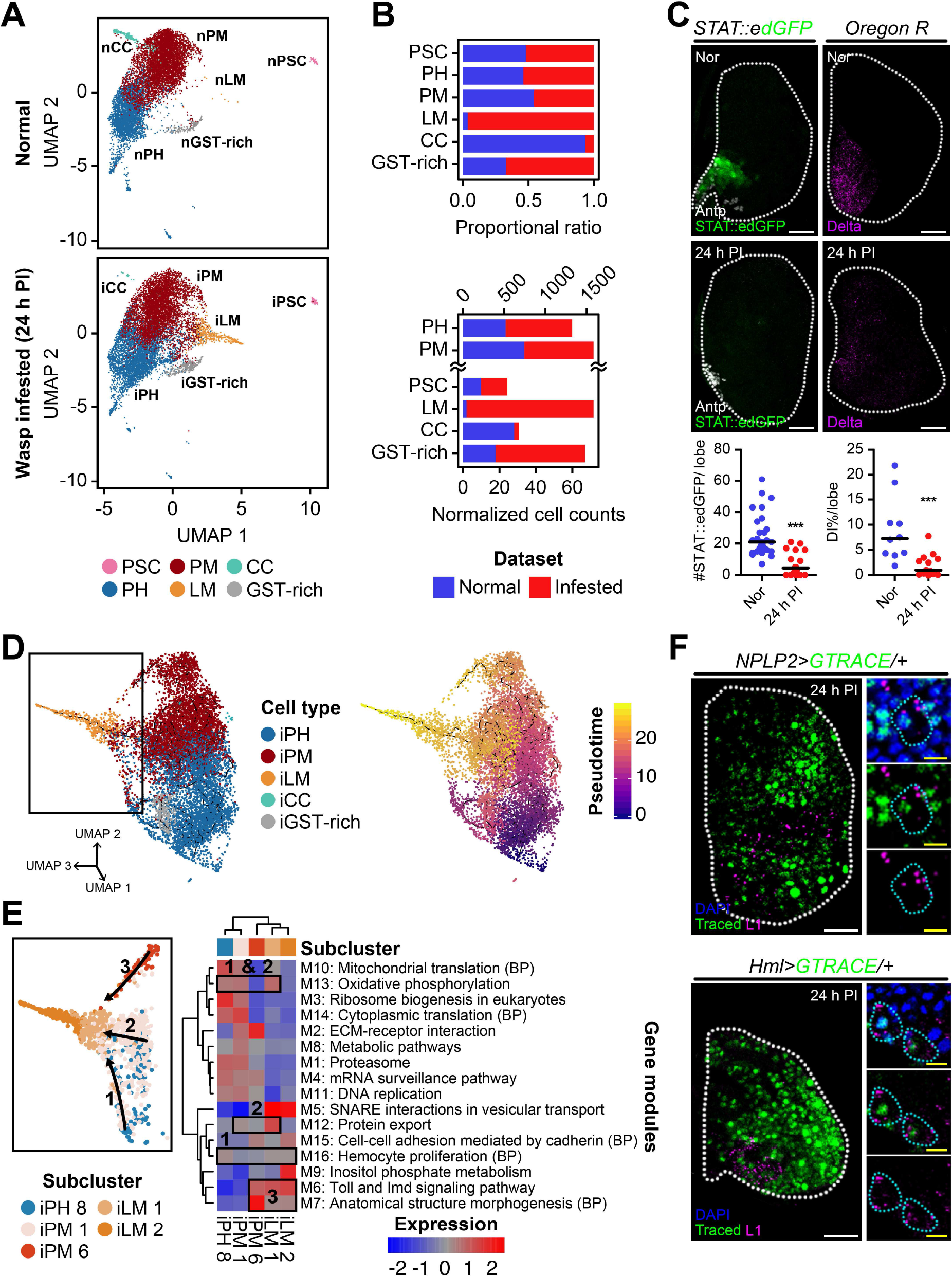
Lymph gland hematopoiesis following wasp infestation. (A) UMAP projections of major cell types defined in normal (top, prefix with ‘n’) and wasp infested (bottom, prefix with ‘i’) lymph glands at 96 h AEL (thus, 24 h post infestation (PI)). Non-hematopoietic cell types are excluded. Different colors indicate each cell type. (B) Relative proportion (top, proportional ratio) or normalized cell counts (bottom) of major cell types in normal (blue) and wasp infested (red) lymph glands. (C) Wasp infestation reduces *STAT92E^+^* (left) or Dl^+^ (right) PH1 populations. *STAT92E* (green) or Dl (magenta) are expressed near the PSC of the lymph gland (top). The expression is attenuated upon immune challenges (middle). Graphs represent quantitation of the number of *STAT92E^+^* cells (bottom, left) or the proportion of Dl^+^ (bottom, right) cells in one lymph gland lobe after infestation (24 h PI). Median value is shown in graphs (bottom, ***p<0.0001). (D) A three-dimensional trajectory landscape of major cell types under wasp infestation (left), and additional representation of trajectory over calculated pseudotime using Monocle 3 (right). Box indicates the cells used for subtrajectory analysis in (E). Colors in legends show pseudotime (right). (E) Subtrajectory analysis of five subclusters—iPH8, iPM1, iPM6, iLM1, and iLM2—detected in the trajectory to iLM (left). Two different waves, arrow 1/2 (iPH8/iPM1, thus, intermediate cells) and arrow 3 (iPM6), advance towards iLM with distinct gene modules (right). Shared gene modules between iPH or iPM with iLM are indicated in boxes. Colored expression represents the *z*-transformed enrichment level of gene modules. (F) Lamellocytes differentiate from intermediate iPHs (*Nplp2-Gal4 UAS-GTRACE*) or iPMs (*Hml-Gal4 UAS-GTRACE*) upon wasp infestation. *Nplp2^+^* iPHs (green, traced; blue, DAPI; top) or *Hml^+^* iPMs (green, traced; blue, DAPI; bottom) express L1 (magenta) upon wasp infestation. Insets indicate magnified images of L1^+^ cells. Cyan dotted lines within insets demarcate traced iLMs. Lymph glands are demarcated by white dotted lines. White scale bar is 30 μm; yellow scale bar is 3 μm.

To better understand how iLM differentiates in the lymph gland, we performed pseudotime trajectory analysis and examined gene modules of related subtypes. Upon the trajectory analysis, we discovered that iLMs are associated with a wide span of iPHs and iPMs (Figure 5D, Figure S5F-S5G). The majority of iLMs are directly derived from iPH8 and iPM1 (route 1 and 2 in Figure 5E), subclusters indicated as the intermediate cell populations (Figure 5E, Figure S5H). Additionally, an alternative route is generated from iPM6 (route 3 in Figure 5E), the most mature plasmatocyte subcluster at 96 h AEL (Figure 5E, Figure S5H). We validated the data by tracing the IZ or the CZ markers upon wasp infestation and confirmed that L1^+^ iLMs are derived from either *Nplp2*^+^ intermediate hemocytes or *Hml*^+^ plasmatocytes (Figure 5F). However, differentiating crystal cells and lamellocyte lineages are mutually exclusive (Figure S5I). An association of gene-expression modules of each subcluster indicates the existence of two distinct trajectories to iLMs: iPH8/iPM1-to-iLM and iPM6-to-iLM (Figure 5E, right). The first iPH8/iPM1-to-iLM wave is enriched with genes involved in hemocyte proliferation and oxidative phosphorylation (Figure 5E, right). The second iPM6-to-iLM wave expresses Toll/Imd pathway and structural genes, demonstrating at least two modes of iLM development in the lymph gland upon wasp infestation. Overall, we established that the lymph gland responds to wasp infestation by an expansion of differentiating hemocytes accompanied by subsequent differentiation of iLMs. In addition to the differentiation of intermediate populations indicated as iPH8/iPM1-to-iLM in the trajectory, mature plasmatocytes, iPM6, trans-differentiate into iLMs as an alternative route amplifying the magnitude of iLM formation.

### Genetic comparison between two hematopoietic lineages

*Drosophila* hematopoiesis occurs in two waves, and hemocytes originating from these two lineages differentiate into indistinguishable cell types (Bazzi et al., 2018; Ghosh et al., 2015; Sanchez Bosch et al., 2019). To distinguish and compare these two lineages, we compared the larval circulating hemocyte dataset (see accompanying paper, Tattikota *et al*.) to explore lineage-specific features of *Drosophila* hemocytes at 96 and 120 h AEL. We performed Seurat alignment to cluster datasets after adjusting for batch effect. We also excluded genes related to stress responses that may have been induced during sample preparation (see Methods for details). As a result, we found that hemocytes from the lymph gland significantly overlap with those from circulation (Figure 6A, Figure S6A). We then transferred subcluster labels of lymph gland hemocytes to circulating hemocytes, and recognized three common cell types: prohemocytes, plasmatocytes, and crystal cells, all of which consisted of seven subclusters in circulation (Figure 6B, Figure S6B). Lamellocytes, adipohemocytes, and the PSC are exclusively found in the lymph gland (Figure 6B). All 67 prohemocytes in circulation are labeled as PH1 with unique markers (Figure S6B-S6C), albeit in the absence of Notch and its downstream components (Figure 6C, Figure S5C).

**Figure 6.**
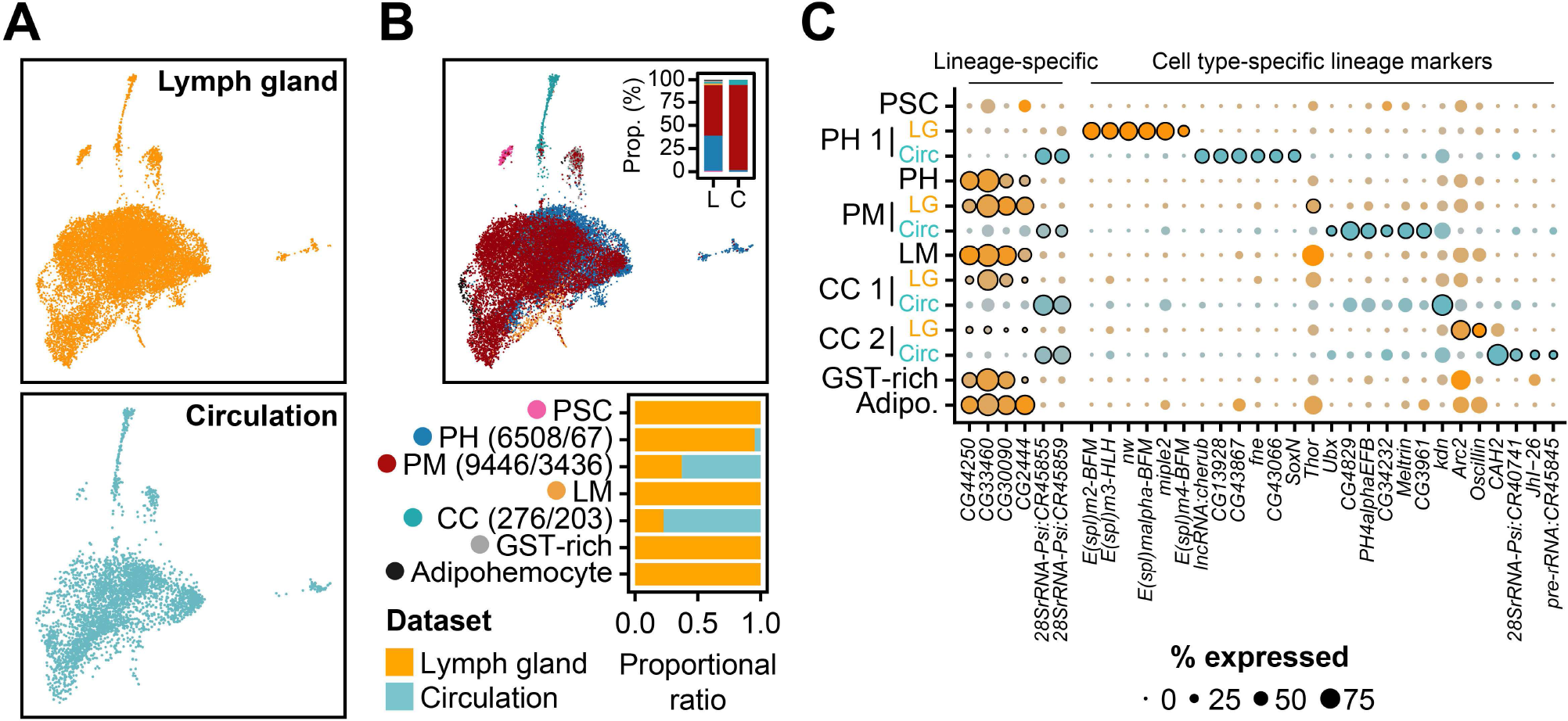
Transcriptome-wide comparisons between embryonic and definitive hemocytes in *Drosophila*. (A) Two-dimensional projections of hemocytes in the lymph gland (top) and circulation (bottom) at 96 and 120 h AEL. (B) Combined projection (top) and proportions of major cell types (bottom) in the lymph gland (yellow) and in circulation (cyan). Inset (top) shows the ratio of major cell types between the lymph gland (L) and in circulation (C). (C) Dot plot of marker genes highly enriched in a lineage-specific or cell type-specific manner. The colors show the origin of the datasets (yellow, lymph gland; cyan, circulation).

Plasmatocytes in circulation share similarities to PM4, PM5, PM6, and PM7 of the lymph gland (Figure S5D). Additionally, crystal cells in the lymph gland and in circulation are nearly identical except for a few genes (Figure S5E-S5F).

Next, we explored the collective signature gene expression of lymph gland and circulating hemocytes. Besides the genes depicted caused by uneven proportions (Figure S6A), we identified novel lineage-specific genes including *28SrRNA-Psi:CR45855* and *28SrRNA-Psi:CR45859* in circulation and *CG44250* and *CG33460* in the lymph gland (Figure 6C; Figure S6A). When each subcluster was individually compared, circulating plasmatocytes display *Ubx* expression while plasmatocytes in the lymph gland show *Thor* expression (Figure S5D, S5G-S5I). Similar differences are observed in crystal cells: *Pde1c*, *CAH2*, and *Naxd* are higher in circulating crystal cells whereas *Arc2*, *Oscillin*, *aay*, and *fbp* are significant in crystal cells from the lymph gland (Figure S5E-S5F). Taken together, hemocytes generated from the two independent lineages appear to be predominantly similar; however, they are sufficiently genetically distinct that we can distinguish their ancestries.

### Evolutionary conservation of lymph gland hemocytes

Although functional homologies between *Drosophila* hemocytes and vertebrate immune cells have been addressed previously (Cooper, 1976; Evans et al., 2003; Gold and Bruckner, 2014), no system-level comparison has been reported. Thus, we compared the single-cell transcriptome profiles of six hematopoietic *Drosophila* cell types including prohemocytes, plasmatocytes, crystal cells, lamellocytes, GST-rich, and adipohemocytes, with human hematopoietic lineages from the Human Cell Atlas (HCA) project (Census of Immune Cells)(Rozenblatt-Rosen et al., 2017). 262,638 high-quality immune cells were clustered and annotated with 19 well-known immune cell types (Figure 7A-7B). Following clustering, we observed a clear separation of lymphoid cells, including T cells and B cells, from the others, while erythroblasts, B-cell precursors, and granulocyte progenitors are closely linked to hematopoietic stem cells∼multipotent progenitor (HSC∼MPP) cluster (Figure 7A-7B).

**Figure 7.**
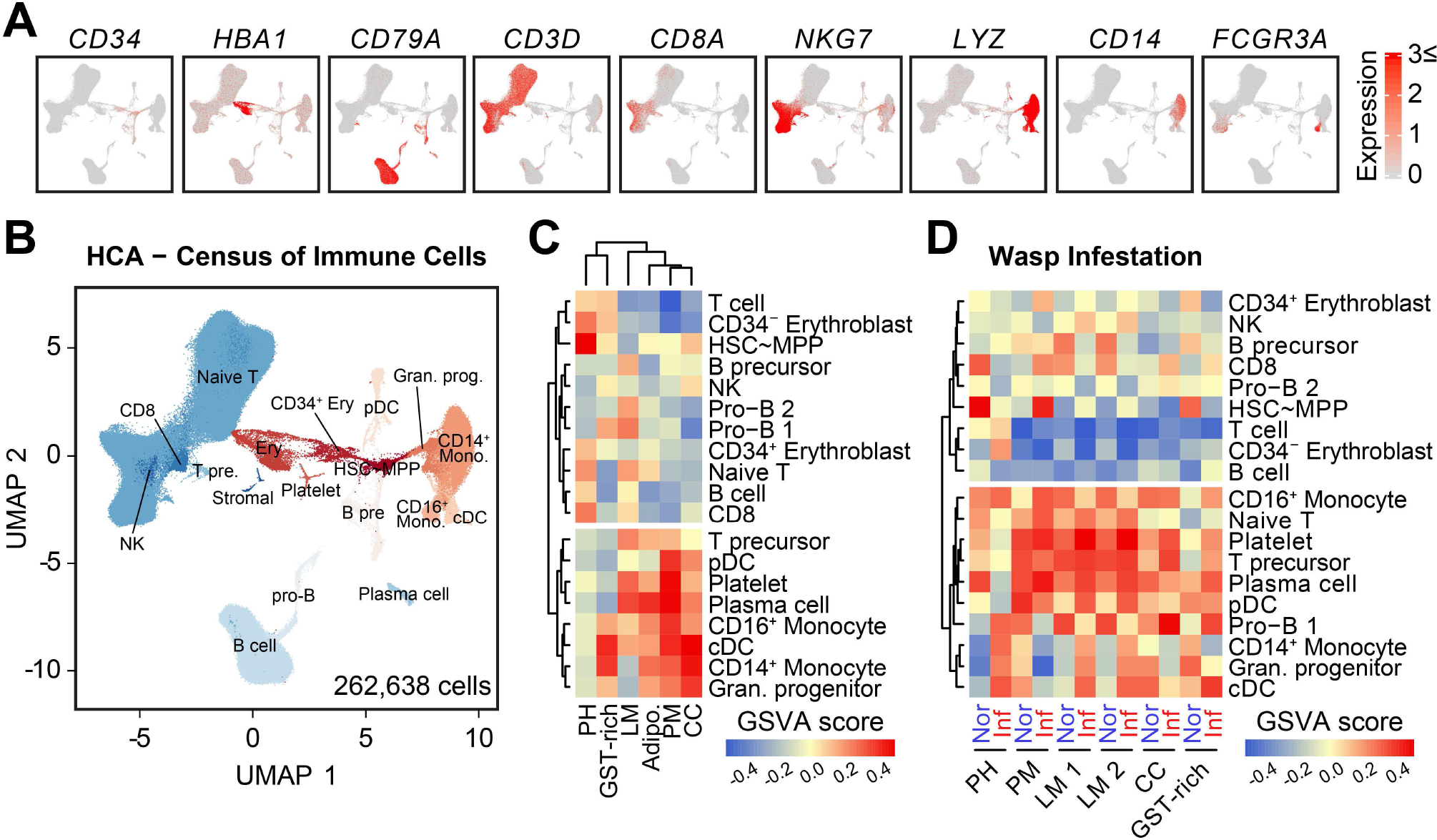
Transcriptome-wide comparisons of *Drosophila* hemocytes and human immune cells. (A) Expression of known markers for human hematopoietic cells in the bone marrow. The colored bar indicates the level of scaled gene expression. CD34 indicates hematopoietic stem cell; HBA1, erythrocyte; CD79A, B cell; CD3D, naïve T cell; CD8A, CD8 T cell; NKG7, NK cell; LYZ, granulocyte progenitor; CD14, CD14^+^ monocyte; and FCGR3A, CD16^+^ monocyte. (B) Annotations of 19 human hematopoietic cell clusters of 262,638 cells provided by the Human Cell Atlas project. (C) Gene set variance analysis (GSVA) of *Drosophila* lymph gland hemocyte and human hematopoietic cell clusters. The colored bar indicates the Gene set variance analysis (GSVA) score. (D) GSVA of hemocytes with human hematopoietic cells from normal (red) and 24 h post wasp infestation (blue) lymph glands at 96 h AEL. The colored bar indicates the GSVA score.

To compare the similarities of gene expression patterns between human immune cells and *Drosophila* hemocytes, 6,463 orthologous gene pairs were retrieved, and enrichment of the top 30 signature genes of each hemocyte type was weighed in human cell types using gene set variance analysis (GSVA) (see Methods for details). Strikingly, we observed a clear separation into two lineages, with one cluster consisting of lymphoid-lineage human cells devoid of *Drosophila* hemocytes, whereas the second one was enriched with myeloid lineage containing most of the hemocytes (Figure 7C; Table S4). Prohemocytes in the *Drosophila* lymph gland share similar gene expression with the HSC∼MPP cluster in humans, while mature hemocytes in general are comparable to human myeloid immune cells (Figure 7C). Specifically, plasmatocytes exhibit similar expression with monocytes, dendritic cells, and platelets, whereas adipohemocytes resemble more plasma cells (Figure 7C). Both crystal cells and GST-rich are closely associated with CD14^+^ monocytes and classical dendritic cells (cDC); however, crystal cells are much closer to granulocyte progenitors than other cell types (Figure 7C). Intriguingly, genetic similarities of *Drosophila* hemocytes and human myeloid cells are extensively affected when immune-challenged hemocytes and human immune cells are compared. Plasmatocytes become comparable to HSC∼MPP, and, surprisingly, *Drosophila* hemocytes at 24 h post-infestation display striking similarities with lymphoid precursor lineages such as naïve T cells, T cell precursors, and pro-B cells of humans, implying the functional duality of active hemocytes beyond their homologies to myeloid cells (Figure 7D). In conclusion, our analysis demonstrates that *Drosophila* hemocytes retain conserved genetic characteristics of broad classes of human myeloid immune cells and are plastic enough to show lymphoid-like features upon active immunity.

## DISCUSSION

In this study, we report a comprehensive single-cell transcriptome analysis of 29,618 developing myeloid hemocytes in *Drosophila* lymph glands. Our analysis provides insights into: 1) the development of myeloid hemocytes at the single-cell level, 2) the existence of hematopoietic stem-like populations and adipohemocytes in invertebrates, 3) the differentiation mechanisms of myeloid hemocytes upon active immunity, 4) the genetic difference of hemocytes derived from two different hematopoietic ancestries: embryo and larva, and 5) the evolutionary relevance of *Drosophila* hemocytes. To our knowledge, this study is the first description of invertebrate myeloid cells at a single-cell level and a system-level comparison of myeloid cells across species.

### scRNA-seq reveals novel hemocyte types in the lymph gland

Unlike human hematopoietic systems, *Drosophila* hematopoiesis takes place in a unique hematopoietic organ, the lymph gland, where all developing myeloid cells are located until histolysis upon pupariation (Grigorian et al., 2011). Thus, scRNA-seq of the lymph gland allows us to analyze inclusive profiles of myeloid hemocytes containing all developmental stages of myeloid cells. Our scRNA-seq datasets faithfully display single-cell transcriptomes of all known cell types as well as two novel cell types: GST-rich and adipohemocytes. GST-rich cells, enriched with ROS-responsive and DNA damage genes, emerge during prohemocyte development. Considering that genes enriched in GST-rich cells are also evident in the lymph gland bulk RNA-seq, this novel population is not a consequence of stressed hemocytes. Rather, this subtype may represent a state that prohemocytes experience during development, or may play an active role in ROS- or GABA-mediated stress responses (Madhwal et al., 2019; Owusu-Ansah and Banerjee, 2009; Shim et al., 2013). Adipohemocytes, on the other hand, share hallmarks of both mature plasmatocytes and lipid metabolism, appearing only at 120 h AEL of the lymph gland. Macrophages in vertebrates readily take up lipids and lipoproteins, and accumulation of lipid-containing macrophages, called foam cells, is highlighted in various pathological conditions (Li and Glass, 2002; Moore et al., 2013). In *Drosophila*, the presence of lipid-containing hemocytes has not been reported. Given our analyses, and that adipohemocytes are frequently observed in insects, including *Aedes aegypti* (Hillyer et al., 2003), it is possible that flies also conserve metabolism-oriented hemocytes to coordinate immunity and metabolism.

### Prohemocytes are comprised of heterogeneous cell types and states

Prohemocytes have been widely considered to represent a uniform cell population based on the expression of marker genes, *domeless* or *Tep4*. However, recent studies have suggested that prohemocytes may be more heterogenous based on uneven expressions of cell cycle markers or bifurcated *col* expressions (Baldeosingh et al., 2018; Sharma et al., 2019). In support of these studies, our unbiased subclustering of primary clusters identified different status of prohemocytes. First, prohemocytes differ in the expression of cell cycle regulators, implying an asynchrony of prohemocyte development and their states. This observation also accounts for the stochastic cell cycle patterns visualized with the UAS-FUCCI system, a fluorescent-based cell cycle indicator (Sharma et al., 2019; Zielke and Edgar, 2015). Second, we observed dynamic expression patterns of immunogenic, metabolic or stress-responsive genes in PH subclusters. For example, PH7 and PH10 are immunogenic while PH4, PH5, PH6, PH8, and GST-rich are metabolic or stress-responsive. These multiple states could be due to different susceptibility of prohemocytes to physiological conditions and may directly influence lineage specification. Lastly, the presence of prohemocytes with more differentiated states is also indicative of their dynamics. Although the presence of the intermediate zone has been recognized in previous studies (Krzemien et al., 2010; Owusu-Ansah and Banerjee, 2009), the biological significance of various intermediary states and the novel functions of expressed endogenous genes including *Nplp2* in these subclusters have not been explored.

### PH1 is the stem-like prohemocyte in the lymph gland

As the most primitive subcluster identified in this study, PH1, demarcates a group of cells that has not been annotated by previous markers such as *Tep4*, *Antp* or *col*. Discovery of the hidden cell population – PH1, will shed light on understanding the hierarchy of prohemocyte differentiation and enhance the relevance of the lymph gland as a hematopoietic model. Roles for *Notch*, *Stat92E*, or *scalloped* in the lymph gland development have been previously suggested by recent studies (Dey et al., 2016; Ferguson and Martinez-Agosto, 2017; Krzemien et al., 2007; Mondal et al., 2011). Moreover, clonal analyses have shown that cells adjacent to the PSC generate the largest population in the lymph gland (Minakhina and Steward, 2010a). These studies are consistent with our hypothesis that Notch/Delta- and JAK/STAT-positive cells nearby the PSC sustain latent capacities to produce the entire lymph gland hemocytes. We also observed that PSC is required for the expression of PH1, and thus, PSC is likely to provide necessary signals for the maintenance of PH1 during normal development. Given that *col*^+^ hemocytes, referred as PH2 in this study, are independent of the PSC (Baldeosingh et al., 2018; Benmimoun et al., 2015), it is possible that there are multiple niches for respective subtypes, such as the PSC for PH1 and the dorsal vessel for PH2/PH3, which are reminiscent of context-dependent niches found in vertebrates (Tikhonova et al., 2019). Identification of the factors underlying the maintenance and differentiation of PH1 will be important for understanding the nature of hematopoietic stem-like populations.

### Hemocytes conserve genetic homologies with human immune cells

We attempted to uncover cellular heterogeneity of identical cell types originating from two different lineages and genetic homologies between *Drosophila* hemocytes and human immune cells. Previously, *Drosophila* hemocytes have been proposed to be most akin to macrophages of vertebrates (Franc et al., 1996; Sanchez Bosch et al., 2019); however, our analysis indicates that *Drosophila* hemocytes show characteristics of multiple human myeloid cells, including monocytes, dendritic cells, and granulocyte progenitors. Furthermore, our analysis reveals that *Drosophila* hemocytes additionally acquire signatures of lymphoid lineages upon active immunity. In light of our analyses, we propose that invertebrate myeloid cells carry hidden elements of lymphoid activity, contributing to divergence of the lymphoid lineage in vertebrates. Collectively, our comparative analysis supports genetic similarities between myeloid cells of flies and vertebrates, providing a resource to further understand the invertebrate and vertebrate myeloid cells.

## Acknowledgements

The authors thank Dr. Greg S. Suh and all members of the Shim and the BIG labs for helpful discussions. The authors acknowledge the Bloomington, VDRC, DGRC, NIG and KDRC *Drosophila* stock centers and the DSHB hybridoma bank. The authors thank the following individuals for stocks and reagents: Drs. C. Evans, U. Banerjee, S. Sinenko, M. Zeidler, M. Crozatier, K. Brueckner, Nambu JR, A. Brand, and F. Schweisguth. This work was supported by the Samsung Science and Technology Foundation under Project Number SSTF-BA1701-15 to J.S. and by the National Research Foundation (NRF) funded by the Ministry of Science and ICT under Project Numbers 2017M3A9G8084539 and 2018R1A2B2003782 to J.N., and 2016R1A5A2008630 to S.J.M. N.P. is an Investigator of the Howard Hughes Medical Institute.

## Author Contributions

B.C., S.G.T., D.L., F.K., N.C., M.S., H.D., and S.Y.O. performed experiments; B.C., S.Y., S.G.T., Y.H., J.N., and J.S. analyzed data; S.Y.O. and S.J.M. provided technical support for Drop-seq; B.C., S.Y., F.K., S.G.T., N.P., J.N., and J.S. contributed to writing the manuscript; N.P., J.N., and J.S. supervised the project; J.N., and J.S. conceived the idea.

## Declaration of Interest

The authors declare no competing interests.

## Material and method

### *Drosophila* stocks and genetics

The following *Drosophila* stocks were used in this study: *Dome^Meso^-EBFP2* (U.Banerjee), *HmlΔ-Gal4* (S.Sinenko), *Dome^Meso^-Gal4* (M.Zeidler), *TepIV-Gal4* (NIG, Japan), *Nplp2-Gal4* (KDRC, South Korea), *pCol85-Gal4* (M.Crozatier), *Delta-Gal4* (Bloomington), *Antp-Gal4* (U.Banerjee), *Stat92E::edGFP* (N.Perrimon), *Hml-DsRed* (K.Brueckner), *vir-1^MiMiC^* (Bloomington), *Ance^MiMiC^* (Bloomington), *UAS-hid, rpr* (Nambu JR), *UAS-GTRACE* (C.Evans), *nSyb-Gal4* (Bloomington), *Ama-Gal4* (NIG), *crq-Gal4* (Bloomington), *Lsd2-Gal4* (NIG), *Ilp6-Gal4* (A.Brand), *Tau^MiMiC^ (*Bloomington), *mthl7-Gal4* (generated in this study), *Chrb^MiMiC^* (Bloomington), *Men-Gal4 (*NIG, 113708), *NUMB::GFP* (F.Schweisguth), *zfh1-Gal4 (*Bloomington), *Tep2^MiMiC^*(Bloomington), *UAS-EGFP* (Bloomington), *UAS-mCD8GFP* (Bloomington), *Ubx RNAi*(v37823), *lz-gal4^DBD^; Pxn-Gal4^AD^* (generated in this study)

Fly stocks used in this study were maintained at 25 °C. *Oregon R* was used for the scRNA-seq as a wild type. Unless indicated, crossed flies were maintained at 25 °C with dextrose-cornmeal based normal food. To synchronize larval stages, one hundred adult flies were kept on grape-juice agar plate for two hours. Hatched larvae were discarded at 23 hours after egg laying (AEL), and those at 24 hours AEL were collected and reared on normal corn-meal yeast media. To screen the Gal4 lines in this study, we crossed each Gal4 strain with *UAS-GTRACE* and identified those expressed in the lymph gland. To avoid stress conditions generated by crowding, we reared less than 50 larvae in one vial.

### *Generation* of fly stocks

To generate Gal4 fly lines, fly genomic DNA was amplified by primers indicated in Supplementary table 3. Amplified genomic regions were ligated into *pAGal4+*(KDRC) or TOPO-TA vector (Invitrogen; K250020) for Gateway cloning. *pBPnlsLexAp65Uw* (Addgene 26230), *pBPZpGAL4DBDUw* (Addgene; 26233) or *pBPp65ADZpUw* (Addgene; 26234) was used as destination vector. Transgenic flies were generated by KDRC, South Korea.

### Dissociation of lymph glands into single cells

100 to 150 lymph glands were dissected at 72, 96, or 120 h AEL respectively, in Schneider’s medium (Gibco, 21720024). Dorsal vessel, ring gland and posterior lobes were detached from the primary lobe of the lymph gland; only the primary lobes of the lymph glands were used in this study. Primary lobes were kept in 200 μl ice-cooled Schneider’s medium during dissection.

After, centrifugation at 3,000 rpm and 4 ℃ for 5 minutes was done. Supernatant was discarded and 300 μl of room temperature Schneider’s medium was added to the lymph gland primary lobes. 300 μl of Papain (Worthington, LK003178) pre-heated to 37 ℃, and 4.1 μl of Liberase TM (Roche, 5401119001) were added and gently mixed. Samples were incubated for 20 minutes with gentle agitation.

At 5-, 10-, and 15-minute time points of incubation, samples were mixed using 200p pipette. Enzymes were inactivated with 100 μl of ice-cooled Schneider’s medium, and samples were kept on ice. Suspended cells were passed through a 40 μm cell strainer (Corning, 352340). Afterwards, centrifugation at 3,000 rpm and 4 ℃ for 5 minutes was done. The supernatant was discarded, and 1X filtered sterile PBS was added to cells. The final concentration of cells was fixed to 300 cells/μl for a total of 600 μl.

### Drop-seq and scRNA-seq

All the Drop-seq and cDNA synthesis methods followed a previous study (Macosko et al., 2015). The concentration of beads was fixed to 300 beads/μl. Around 10 minutes Drop-seq run was performed for each experiment. After cDNA synthesis, scRNA-seq was performed using Illumina NextSeq.

### Preprocessing and mapping of scRNA-seq data

Raw scRNA-seq data were generated in paired-end reads following single-cell capture using Drop-seq: one end included a barcode and unique molecular identifier (UMI) sequences in 20 nucleotides (12 and 8 nts, respectively), and the other end, cDNA in 50 nts. The preprocessing and mapping of scRNA-seq data produced in this study followed the Drop-seq Core Computational Protocol version 1.2 (January 2016) and corresponding Drop-seq tools version 1.13 (December 2017) provided by the McCarroll Lab (http://mccarrolllab.org/dropseq/).

First, the reference genome (Fasta, BDGP 6.02) and transcriptome annotations (gtf, September 2014) required for the processing were downloaded from the Ensembl website (http://asia.ensembl.org/). Additional dictionary and refFlat files were generated using *picard* (*CreateSequenceDictionary*) and *ConvertToRefFlat* provided in the Drop-seq tools package, respectively. These reference data were prepared with the same prefix and stored in a single directory for later use. Simplified command lines are as follows:

Generation of a dictionary file:

~~~
*java -jar <PATH to Drop-seq tools picard>/picard.jar
CreateSequenceDictionary R=GENOME fasta> O=<OUTPUT dictionary>*
~~~

Generation of a refFlat file:

~~~
*<PATH to Drop-seq tools>/ConvertToRefFlat \
ANNOTATIONS_FILE=<GFT annotation> \
SEQUENCE_DICTIONARY=<DICTIONARY file> O=<OUTPUT refFlat>*
~~~

Once all the reference data was prepared, paired-end fastq files were converted to the bam format using *picard FastqToSam*.

~~~
*java -jar <PATH to Drop-seq tools picard>/picard.jar FastqToSam \
F1=<FASTQ 1> F2=<FASTQ 2> O=<OUTPUT bam> SM=<LIBRARY number>*
~~~

The unaligned bam files were subjected to the *Drop-seq_alignment.sh* script for alignment to genome. This shell script is a single pipeline that executes detection of barcode and UMI sequences, filtration and trimming of low-quality bases and adaptors or poly-A tails, and alignment of reads using *STAR* (2.5.3a).

~~~
*<PATH to Drop-seq tools>/Drop-seq_alignment.sh \
-g <PATH to STAR index> -r <GENOME fasta> -n <# of cells expected> \
-d <PATH to Drop-seq tools> -s <PATH to STAR> \
-o <PATH to output> -t <PATH to temporary output> -p <UNALIGNED bam file>*
~~~

### Selection of cells by the total mapped reads

To extract the number of cells having proper read counts, the aligned bam files generated from the previous section were summarized using *BAMTagHistogram* in the Drop-seq tools package. This program extracts the number of aligned reads per cell barcode which is subsequently used to plot the cumulative distribution of reads.

~~~
*<PATH to Drop-seq tools>/BAMTagHistogram I=<ALIGNED bam> O=<OUTPUT file> TAG=XC*
~~~

Cumulative read distribution plots were then explored, and the number of cells were inferred where a sharp decrease (referred as ‘knee’ by the author’s documentation) in a slope occurs. The inferred cell number was determined as a minimal threshold number of aligned reads per cell for cell selection. To summarize, a minimum of 30,000 reads per cell for 72 h AEL library 6, 15,000 per cell for 72 h AEL library 3, 10,000 per cell for 72 h AEL libraries 1, 2, and 4; 5000 per cell for 96 h AEL libraries 1, 3, 5, all 120 h AEL libraries, and infested 96 h AEL libraries 1, 2, and 4; 4000 per cell for infested 96 h AEL 2, 3000 per cell for 96 h AEL library 4 and infested 96 h AEL library 3; 2000 per cell for 92 h AEL library 2 were chosen as thresholds. *DigitalExpression* provides UMI count matrix (selected cells by genes) using a mapped bam file and the minimum number of reads per cell as following.

~~~
*<PATH to Drop-seq tools>/DigitalExpression \
I=<ALIGNED bam> MIN_NUM_READS_PER_CELL=<READ count threshold> \
O=<OUTPUT read count matrix> SUMMARY=<OUTPUT summary>*
~~~

The resulting output per library is written into a file with a name where a corresponding library number was added as a suffix to each barcode sequence with an AEL timepoint (*e.g.* barcode-72-1 or barcode-96-2) to avoid collision of barcode sequences between libraries. The number of expressed genes between libraries or sampling timepoints may vary because each library would have a different number of captured cells, different cell types, or uneven sequencing depth. In total, at least 13,612, 13,523, 14,277, and 13,658 genes, and 2505, 10,027, 11,702, and 10,939 cells were detected in one library at 72, 96, 120, or infested 96 h AEL respectively. So, we used a union set of genes (15,540 genes) to merge three normal lymph gland datasets.

### scRNA-seq data analysis using Seurat 3.0

Seurat is a universal software for scRNA-seq analyses including preprocessing, cell clustering, and dimension reductions. The current version (v3.0) of Seurat features dimension reduction using uniform manifold approximation and projection (UMAP), integration of datasets produced with different modalities or conditions, and transfer of cell labels between datasets (Butler et al., 2018; Stuart et al., 2019). Detailed analyses steps are explained on the Seurat website (https://satijalab.org/seurat/), so we only describe the schematic workflow used in this study.

First, each library was filtered for low-quality cells, separately, by setting thresholds for UMI and gene counts. We used 5,000 genes as an upper threshold and 400 genes as a lower threshold for normal lymph gland libraries and infested lymph gland library 1 and 2, and 200 genes as a lower threshold for other infested lymph gland libraries. Then we also filtered cells having UMI counts higher than two standard deviations from the mean UMI count to exclude multiplets. After filtration, 2399, 9496, 11,081, and 10,461 cells remained for 72, 96, 120, and infested 96 h AEL, respectively. All libraries in each sampling timepoint (h AEL) were then merged and normalized, and cells expressing mitochondrial genes higher than 10% of total UMI count were removed (2322, 9411, 10,976, and 10,179 cells remained). After high mitochondrial cells were excluded, multiplets were inferred using Scrublet (Wolock et al., 2019), which simulates artificial doublets using given expression matrix to predict multiplet artifacts. One, 11, and 52 cells were further filtered out from 72, 96, and 120 h AEL lymph glands. However, no cells were detected from the infested dataset. As a result, 2321 (72 h AEL), 9400 (96 h AEL), 10,924 (120 h AEL), and 10,179 cells (infested 96 h AEL) were subjected to downstream analyses.

Next, for the integration of cells from normal lymph glands at three timepoints (normal lymph gland integration), cells were aligned using *FindIntegrationAnchors()* and *IntegrateData()* with default parameters, respectively. UMI counts were normalized, log-transformed, and scaled to properly integrate datasets, and 52 principle components (PCs) were used to explain the variability of the scaled UMI counts across cells. t-SNE and UMAP plots were then manually curated with random seeds using the selected PCs. Clustering was performed with resolution of 0.8 to get 19 clusters. Then again, clusters were aggregated to get broad cell types based on expression of known marker genes (Figure S1G). In summary, six and seven clusters were merged as collective PH and PM, respectively, to define the following six major cell types: PSC, LM, CC, DV, GST-rich, and adipohemocyte.

When the integrated normal lymph gland dataset was examined, we found that subclusters of PH and PM cells solely originated from 120 h AEL (Figure 1D and Figure S1I). Thus, we analyzed our dataset using a different batch normalization method to test whether this trend is independent of our analysis strategy. For this, we corrected the sequencing library variable using *ScaleData* function along with the UMI count. We then selected the number of PCs to use (50 PCs) and performed t-SNE and UMAP analyses again. Similarly, a number of cells from 120 h AEL were separated from others while cell types, such as PSC, LM, or CC were mixed together (Figure S1J). An R package, *rgl*, was used for all 3-dimensional plots presented in this study.

### Pseudo-bulk RNA-seq trajectory analysis

Normal lymph gland scRNA-seq libraries were examined to see whether they could be aligned in a single trajectory line as actual development timepoints. As we produced at least four sequencing libraries for each timepoint from independent sample preparations and sequencing, we performed trajectory analysis in pseudo-bulk RNA-seq samples, pooling all cells from each library. For this, valid cell barcodes identified in the previous analysis were collected and their UMI count matrices were retrieved. All UMI count values were then aggregated by genes to generate pseudo-bulk RNA-seq data. We applied Monocle 2 (Qiu et al., 2017; Trapnell et al., 2014) for this trajectory analysis using 2758 highly variable genes or top 500 most differentially expressed (DE) genes out of 7596 genes (expressing more than one UMI in at least three cells) with default parameters. The highly variable genes were selected with criteria of *dispersion_empirical >= 1 * dispersion_fit & mean_expression >= 1*. In both analyses, the results were similar, so the trajectory using variable genes was presented (Figure S1E).

### Subclustering analysis

To determine detailed cellular states or subtypes and to exclude unintended cells that originate from only a single library, each of eight major cell types was clustered separately using Seurat. For this, 1) cells designated to each cell type were retrieved from the integrated Seurat object, 2) their UMI counts were scaled and normalized for sequencing library and total UMI counts, and 3) the number of PCs were determined for dimension reduction and clustering analysis. In each subclustering analysis, hundreds of t-SNE and UMAP plots were manually examined with random seeds for visualization. Resolution for clustering analysis was manually selected as follows: 13 subclusters for PH (*res* = 0.8), 14 for PM (*res* = 0.9), two for LM (*res* = 0.3), three for CC (*res* = 0.1), and another three for adipohemocytes (*res* = 0.3) were detected. Other cell types (PSC, GST-rich, and DV) were not further clustered. Next, subclusters mainly originating from a single library (more than half) were removed from the datasets because they may have resulted from experimental artefacts. In fact, two, three, one, and two subclusters were excluded from PH, PM, CC, and adipohemocyte, correspondingly (Figure S2B).

When the results were examined using known marker genes, one PM subcluster (PM11) that displayed a high level of ring gland marker gene, *phm*, was excluded from subsequent analyses. Particularly, of 11 remaining PH subclusters, one PH subcluster (PH1) displayed a high level of genes related to the Notch signaling pathway, which is known to be important for prohemocyte maintenance and differentiation. The PH1 subcluster forms a small group of cells separated from the main PH cluster in the first clustering analysis, which seems to be more related to the PH2 subcluster displaying distinct molecular signatures (Figure 2B). We thus sought to investigate these subclusters in more detail, along with PSC cells which are known to maintain early prohemocytes in lymph glands. For this, we reclustered PH1, PH2, and PSC cells together, and found 6 subclusters. However, one of these subclusters appeared to be neuronal cells as they expressed specific neuronal markers, such as *nSyb* or *Syt1* (Figure S2D). On the other hand, three PSC subclusters were detected but merged because they all shared the same signature genes (Figure 3D). In addition, 13 PH cells— annotated as PSC and 3 PSC cells—that comingled with PH1 or 2 in the subclustering analysis were removed, as their identities were inconsistent. In summary, we defined a total of 31 subclusters from the normal lymph gland dataset (Figure 2); 11 for PH, 10 for PM, two each for LM and CC, one each for PSC, GST-rich, and adipohemocyte, and three non-hematopoietic cell types (DV, RG, and Neurons).

### Trajectory analysis using Monocle 3

To reconstruct lymph gland hematopoiesis in *Drosophila*, hematopoietic cells in the blood lineage were collected after filtering the PSC, DV, RG, and neuron subclusters. We then followed the Monocle 3 (Cao et al., 2019) analysis pipeline described in the website documentation (https://cole-trapnell-lab.github.io/monocle3/) using custom parameters predetermined from repetitive analyses. We normalized the dataset by log-transformation and size factor correction, following three covariates, sequencing library, UMI count, and mitochondrial gene contents (the proportion of mitochondrial genes in transcriptome), and scaled using the *preprocess_cds()* function with 75 PCs. UMAP dimension reduction (*reduce_dimension*) was performed with custom parameters *umap.min_dist* = 0.4 and *max_components* = 3, and clustering resolution (*cluster_cells*) was set to 0.001 which assigned all cells into a single partition. After graph learning was performed (*learn_graph*), the cells were ordered using *order_cells()* to set a node embedded in the PH1 subcluster as a start point. All the trajectory graphs were visualized using the *plot_cells()* function with or without a trajectory graph.

Monocle 3 offers several approaches for differential expression analyses using regression or graph-autocorrelation. In this study, we identified co-regulated genes along the pseudotime by graph-autocorrelation and modularized them. To detect co-regulated genes, the graph-autocorrelation function *graph_test()* was specified with a “principal_graph” parameter and significant genes were selected (*q* value < 0.05). Modularization was performed using *find_gene_modules()* with default parameters but only passing a list of resolution values from 10^-6^ to 0.1 for automatic parameter selection. A total of 51 gene modules were detected from the normal lymph gland trajectory, however, three modules were excluded in that they were unable to be characterized by the enrichment analysis with biological process gene ontology terms or KEGG pathways using g:Profiler (https://biit.cs.ut.ee/gprofiler/gost).

The trajectory analysis of infested lymph gland (Figure 5D) followed a similar pipeline as that of normal lymph gland with slightly different parameters. The dataset was normalized and corrected for covariates, sequencing library, UMI count, and mitochondrial gene contents in the same manner, though using 50 PCs. We then used 0.5 for the minimum distance in UMAP and 0.005 for the clustering resolution. Gene modules were also explored using a complete dataset and the result generally agreed with the previous gene modules using normal lymph glands. So, we focused on the LM trajectory in the analysis. First, we filtered all the other subclusters except for PH8, PM1, and PM6 with two LM subclusters because these subclusters were mainly found in the LM differentiation paths. Then, we collected cells by excluding those having a UMAP 1 coordination higher than −1.5 (Figure 5E, inset). Modularization of co-regulated genes was performed as previously described using these cells.

### Investigation of transcription factor activity using SCENIC

To predict transcription factor regulatory networks in developing lymph glands, we performed SCENIC (Aibar et al., 2017) analyses on the 1) normal lymph gland dataset excluding non-hematopoietic cells (*n* = 19,332) and 2) early PH subclusters with PSC (PH1, PH2, and PSC; *n* = 77, 79, and 189, respectively). We followed the general SCENIC workflow from gene filtration to binarization of transcription factor (TF) activity described by the authors (https://www.aertslab.org/#scenic), using a provided cisTarget reference based on *Drosophila melanogaster* release 6.02. For the latter analysis, we filtered genes using the following two criteria: 1) genes expressed higher than a UMI count threshold of 3*(2.5% of total cell count), and 2) genes expressed in at least 2.5% of total cells (8625 cells). We retrieved 4588 genes from these steps and performed the analysis.

When we investigated the normal lymph gland dataset, however, predictions for several well known TFs in cell types with relatively small sizes, such as *lz* for CC (1.44% of the input dataset) or *Antp* for PSC (0.97% of the input dataset), were affected by the presence of other major cell types. For example, *lz* was an active TF of CC only when PSC, adipohemoctye and GST-rich cell types were excluded from the analysis. We reasoned that the relative population size may affect the predictive power, and signals of TFs active in small populations would be frequently ignored. To tackle this issue, we randomly sampled 42 cells (two thirds of the smallest subcluster—LM2, *n* = 63) from each of 28 hematopoietic subclusters, then collected active TFs using a SCENIC workflow with slightly different gene filtration parameters (we modified 2.5% of the total cell count to 1.0% of total cell count in both gene filtration criteria). We iteratively performed 100 independent trials measuring frequencies for TFs and collected 45 out of 177 TFs predicted active at least 25 times. Then we generated pseudo-bulk profiles for cell types in each timepoint by aggregating scaled gene expression values retrieved from the Seurat object. During the pseudo-bulk generation, LM, CC, GST-rich, and adipohemocyte from 72 h AEL (*n* = 2, 4, 14, and 1, respectively) and LM and adipohemocyte from 96 h AEL (*n* = 14 and 5, respectively) were filtered because the number of cells in these groups was less than 0.1% (∼19 cells) of the complete lymph gland dataset. Using pseudo-bulk profiles, we produced a heatmap describing TF expression in cell types for each timepoint (Figure S1K) using the R package *pheatmap*.

### Comparative analysis between normal and wasp infested lymph glands

For the integration of infested lymph gland (24 h PI; 96 h AEL) with normal dataset from 96 h AEL, those datasets were aligned using Seurat 3.0. functions, *FindTransferAnchors()* and *TransferData()* with default parameters. Again, UMI counts were normalized, log-transformed, and scaled to properly integrate datasets, and the number of PCs was determined (46 PCs). Subclustering labels defined in normal 96 h AEL lymph gland were transferred to infested lymph gland, and three minor subclusters were excluded because they were only found in the normal dataset; PH11 (*n* = 3), PM10 (*n* = 3), and adipohemocyte (*n* = 5). Three non-hematopoietic cell types (35 DV, 18 neurons, and 12 RG cells) were not considered in subsequent analysis.

### Comparative analysis between lymph gland and circulation

A UMI count matrix of 120 h AEL circulating hemocytes described in Tattikota *et al*. was used. Given that the gene annotation version used in Sudhir *et al*. was different from ours, an FBgn to Annotation ID conversion table was downloaded from FlyBase (https://flybase.org; fbgn_annotation_ID_fb_2019_03.tsv) to match gene IDs. After filtration of non-hematopoietic cell types (DV, RG) from the lymph gland dataset and circulating hemocytes expressing high mitochondrial genes (higher than 20%), 17,122 cells from 96 and 120 h AEL lymph glands (*n* = 9339 and 7783, respectively) were aligned with 3706 hemocytes from Tattikota *et al*. dataset and additional 96 and 120 h AEL circulation datasets (*n* = 1235, 977, and 1494, respectively) using 54 PCs. Subclustering labels in lymph glands were transferred to the circulation datasets using *FindTransferAnchors()* and *TransferData()* with default parameters. Prior to the comparison of aligned cell types, a number of subclusters in circulation datasets were filtered because they were either 1) subclusters mainly originated from a single library (more than half) or 2) minor subclusters (less than 1% of total circulation cells). After filtration, seven subclusters remained for subsequent analysis (PH1 = 67, PM4 = 355, PM5 = 1224, PM6 = 961, PM7 = 896, CC1 = 99, and CC2 = 104). To compare gene expression between circulating and lymph gland hemocytes, each cell type or subcluster was independently analyzed for differentially expressed genes using Seurat. Then, gene expression values from each dataset were averaged and log-transformed and top 10 DEGs were marked for visualization (Figure S6).

### Annotation of HCA bone marrow cells

The preview dataset produced by HCA, Census of Immune Cells (Rozenblatt-Rosen et al., 2017), was downloaded from the HCA data portal (https://preview.data.humancellatlas.org). The dataset was produced from bone marrow samples donated by eight individuals and sequenced using 10X Genomics Chromium. To capture high-quality cells from the raw data, we filtered cells that fail to meet the following criteria: 1) those expressing less than 500 genes, 2) those with UMI counts higher than two standard deviations from mean UMI counts, 3) those expressing mitochondrial genes higher than 10% of total gene expression. After the preprocessing step, 262,638 cells were passed to the subsequent analysis. Cells from different donors were aligned using *FindIntegrationAnchors()* and *IntegrateData()* with default parameters, respectively. UMI counts were normalized, log-transformed, and scaled to properly integrate datasets, and 61 PCs were then used for UMAP projection and clustering. Clustering was performed with resolution of 0.8 to get 34 clusters. All the 34 clusters were annotated based on expression of known marker genes. Clusters were then compared using Spearman correlation of gene expression values, and similar clusters (Spearman’s *rho* ≥ 0.95) were aggregated to obtain the 19 clusters shown in Figure 7A.

### Gene set variance analysis of *Drosophila* hemocyte signature genes

Orthologous genes between *Drosophila* and human were searched using DRSC Integrative Ortholog Prediction Tool (DIOPT) version 8.0 (Hu et al., 2011) with default settings. The gene expression of 262,638 HCA cells was aggregated by annotated cell types to obtain a pseudo-bulk matrix, and the non-hematopoietic stromal cell cluster was excluded. Then, 6463 conserved human genes which met one of the following criteria were converted to those of the fly: 1) pairs of genes conserved in one-to-one manner or 2) pairs of genes scored with the highest weighted DIOPT score when conserved in a many-to-one or one-to-many manner.

Signature genes of seven major hemocytes were analyzed by Seurat DEG analysis using *FindAllMarkers()* with *min.pct = 0.25* and *only.pos = TRUE* paramters and only statistically significant genes were left (adjusted *P*-value ≤ 0.05; false discovery rate). Then, these genes were searched in fly-human gene pairs to count the number of signature genes conserved in each major cell type. PH had the least number of conserved signature genes (*n* = 30), and thus, the top 30 conserved signature genes were used for enrichment analysis in other cell types. Gene set variance analysis (GSVA) (Hanzelmann et al., 2013) was performed on pseudo-bulk HCA data with these customized gene sets to identify the enrichment of hemocyte signatures in human bone marrow. GSVA scores were clustered and visualized using an R package *pheatmap* (Figure 7C).

### Bulk RNA-seq of the lymph gland

Procedures prior to the Papain treatment were performed to collect intact lymph glands. Instead of Papain solution, TRIzol (MRC, TR118) was added to lymph glands and RNA extraction performed. More than 1ug of RNA was prepared for each experiment. A cDNA library was constructed by 5’ and 3’ adapter ligation and loaded into a flow cell where fragments are captured into library adapters. Each fragment was then amplified through bridge amplification. After cluster generation was completed, templates were sequenced by Illumina TruSeq.

### Wasp Infestation

Larvae were infested at 72 h AEL with *Leptopilina boulardi*. Wasps were removed after 8 hours of co-culture and egg deposition was confirmed by direct observation of wasp eggs in the hemolymph during dissection. All infestation procedures were performed at 25 °C.

### Immunohistochemistry

Lymph glands were dissected and stained as previously described (Jung et al., 2005). The following primary antibodies were used in this study: α-lz (DSHB, 1:10), α-Antp (DSHB, 1:10), α-Dl (DSHB, 1:10), α-L1 (I.Ando, 1:100), α-col (M.Crozatier, 1:400), α-Ubx (DSHB, 1:10), α-nc82 (DSHB, 1:10), α-NimC1 (I.Ando, 1:100), α-Pxn (Yoon et al., 2017), α-GFP (Sigma Aldrich; G6539; 1:2000) and α-F-actin (ThermoFisher; A34055, 1:200). Cy3-, FITC- or Alexa Fluor 647-conjugated secondary antibody (Jackson Laboratory) was used for staining at a 1:250 ratio. All samples were kept in VectaShield (Vector Laboratory) and imaged by a Nikon C2 Si-plus confocal microscope.

For α-Delta staining, a pre-absorption step was essential to reduce background in the lymph gland. For the pre-absorption step, 1:10 diluted (with 2% sodium azide) α-Dl antibody was incubated together with 9 fixed larval cuticles overnight at 4°C. Lymph glands were dissected in Schneider’s medium and cultured in Schneider’s medium with 10mM EDTA for 30 minutes. Samples were incubated with 3.7% formaldehyde for 1 hour for fixation at 25 °C. After fixation, lymph glands were washed 3 times in 0.1% Triton X-100 in 1xPBS and blocked in 1% BSA/0.1% Triton X in 1xPBS for 1 hour. Lymph glands were incubated overnight at 4°C with α-Dl antibody. Lymph glands were washed 3 times in 0.1% Triton X in 1xPBS and then incubated with α-mouse secondary antibody with 1% BSA/0.1% Triton X in 1xPBS for 3 hours at room temperature. After washing 3 times with 0.1% Triton X in 1xPBS, samples were mounted in Vectashield (Vector Laboratory) with DAPI and imaged by a Nikon C2 Si-plus confocal microscope.

### Bleeding hemocytes

Larvae were vortexed with glass beads (Sigma G9268) for one minute before bleeding to detach sessile hemocytes (Petraki et al., 2015). Larvae were bled on a slide glass (Immuno-Cell Int.; 61.100.17) and hemocytes allowed to settle onto the slide at 4 °C for 40 minutes. Hemocytes were washed 3 times in 0.4% Triton X-100 in 1x PBS for 10 minutes and blocked in 1% BSA/0.4% TritonX in 1xPBS for 30 minutes. Primary antibody was added and samples incubated overnight at 4°C. Hemocytes were washed 3 times in 0.4% Triton X in 1xPBS and then incubated with a secondary antibody with 1% BSA/0.4% Triton X in 1xPBS for 3 hours at room temperature. After washing 3 times with 0.4% Triton X in 1xPBS, samples were kept in Vectashield (Vector Laboratory) with DAPI and imaged by a Nikon C2 Si-plus confocal microscope.

### Quantification of samples

Stained or fluorescent cells were quantified and analyzed by IMARIS software (Bitplane). Individual primary lobes were counted for this study. *In vivo* data was analyzed by the Wilcoxon rank sum test after determining normality with the use of SPSS (version 24).

### Fluorescent *in situ* hybridization

An *in situ* hybridization protocol in a previous study (Crozatier et al., 1996) was used. An anti-*delta* probe was designed based on *delta* cDNA sequences (Forward primer: ATGTGCGAGGAGAAAGTGCT, Reverse primer: CGACTTGTCCCAGGTGTTTT). DIG-labeled probes were detected by α-DIG-biotin antibody (Jackson Immunoresearch; 200-062-156) and visualization was done using a SuperBoost^TM^ Kit (ThermoFisher; B40933). The sense probe was used as a negative control.

### Data and software availability

In-house R and Python codes that were implemented in this study are available on GitHub (https://github.com/sangho1130/Dmel_Dropseq). Raw scRNA-seq and bulk RNA-seq reads are available through the NCBI Gene Expression Omnibus (GEO) (GSE141275).

