## Supplementary Figure 1, Supplementary Figure 2, Supplementary Figure 3, Supplementary Figure 4, Supplementary Figure 5, Supplementary Figure 6 for "Single-cell transcriptome maps of myeloid blood cell lineages in *Drosophila*"

**Supplementary information**

1. Supplementary Figures
2. Supplementary Tables
3. Supplementary Video

### Supplementary Figures

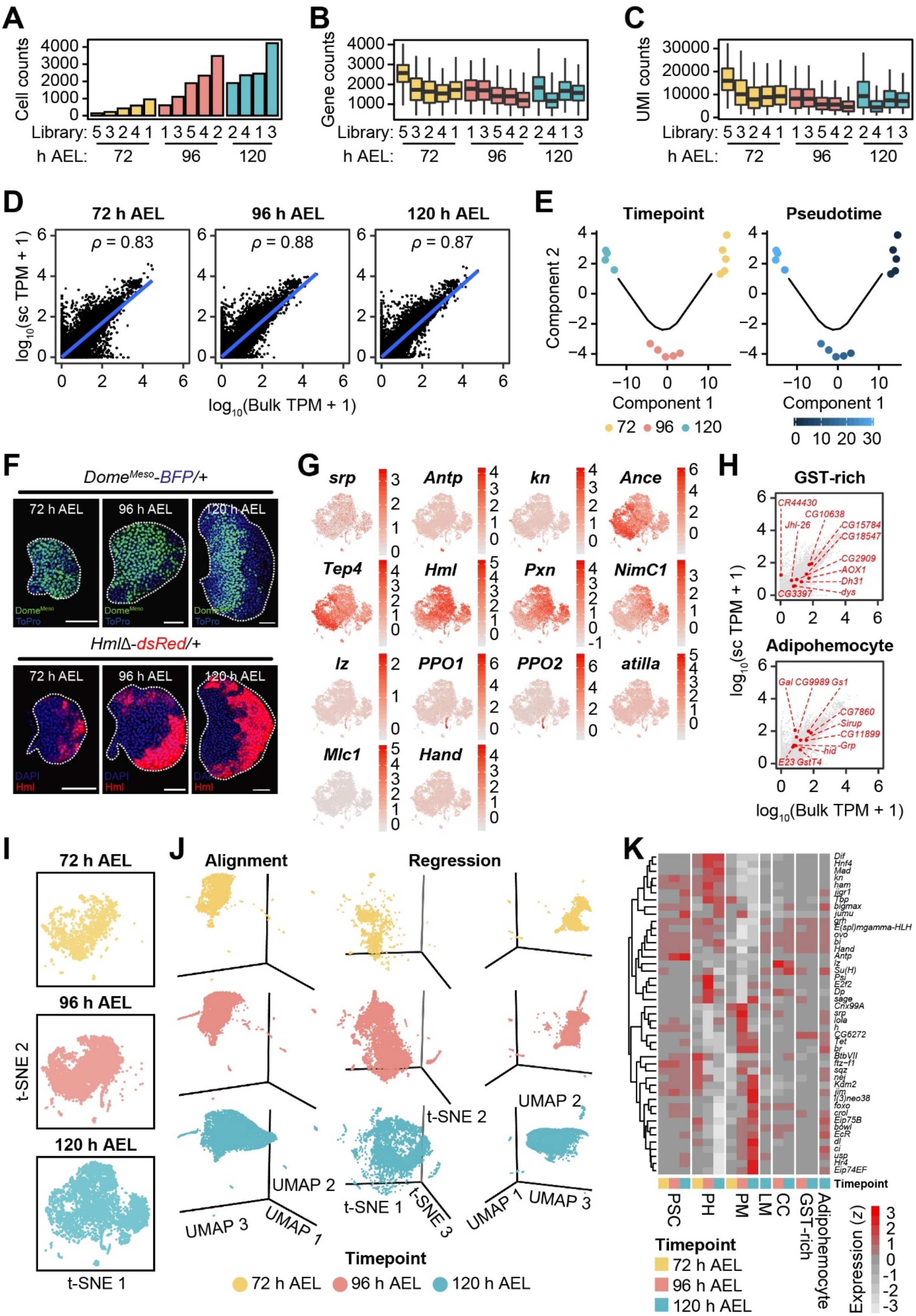

**Figure S1. scRNA-seq of *Drosophila* lymph glands**

(A) Cell counts per each sequencing library are colored by sampling time points. Libraries are ordered by the number of cells (72 h, yellow; 96 h, pink; 120 h, blue).

(B and C) Box plots of gene (B) or UMI (C) counts per sequencing library colored by sampling time points (72 h, yellow; 96 h, pink; 120 h, blue). Libraries are ordered as in (A).

(D) Correlation analyses between pseudo-bulk scRNA-seq and matched bulk RNA-seq samples. Expression values are transformed to the $\log_{10}$ scale.

(E) Trajectory analyses of pseudo-bulk scRNA-seq libraries using Monocle 2 colored by each time point (left) or pseudotime (right) (72 h, yellow; 96 h, pink; 120 h, blue).

(F) Expressions of $Dome^{Meso}$-positive prohemocytes (green, $Dome^{Meso}$; blue, DAPI) or *Hml*-positive plasmatocytes (red, *Hml*; blue, DAPI) at 72, 96, or 120h after egg laying (AEL). White scale bar indicates 30μm. Lymph glands are demarcated by white dotted lines.

(G) Expressions of known marker genes in the major cell types. Color bar indicates the level of scaled gene expression.

(H) Expressions of top 10 signature genes of GST-rich (top) or adipohemocyte (bottom) in pseudo-bulk scRNA-seq and matched bulk RNA-seq. Expression values from three time points in each dataset were averaged and transformed to the $\log_{10}$ scale.

(I) Separation of integrated *t*-SNE projection (related to Figure 1C) of wild-type lymph glands by time points.

(J) Three-dimensional UMAP projection of aligned hemocytes from the lymph gland (left). Three-dimensional *t*-SNE (middle) and UMAP (right) plots of lymph gland hemocytes normalized using regression.

(K) Heatmap presentation of transcription factors in the lymph gland analyzed by SCENIC. Annotations at the bottom indicate time points and major cell types (72 h, yellow; 96 h, pink; 120 h, blue). Color bar indicates the level of scaled gene expression.

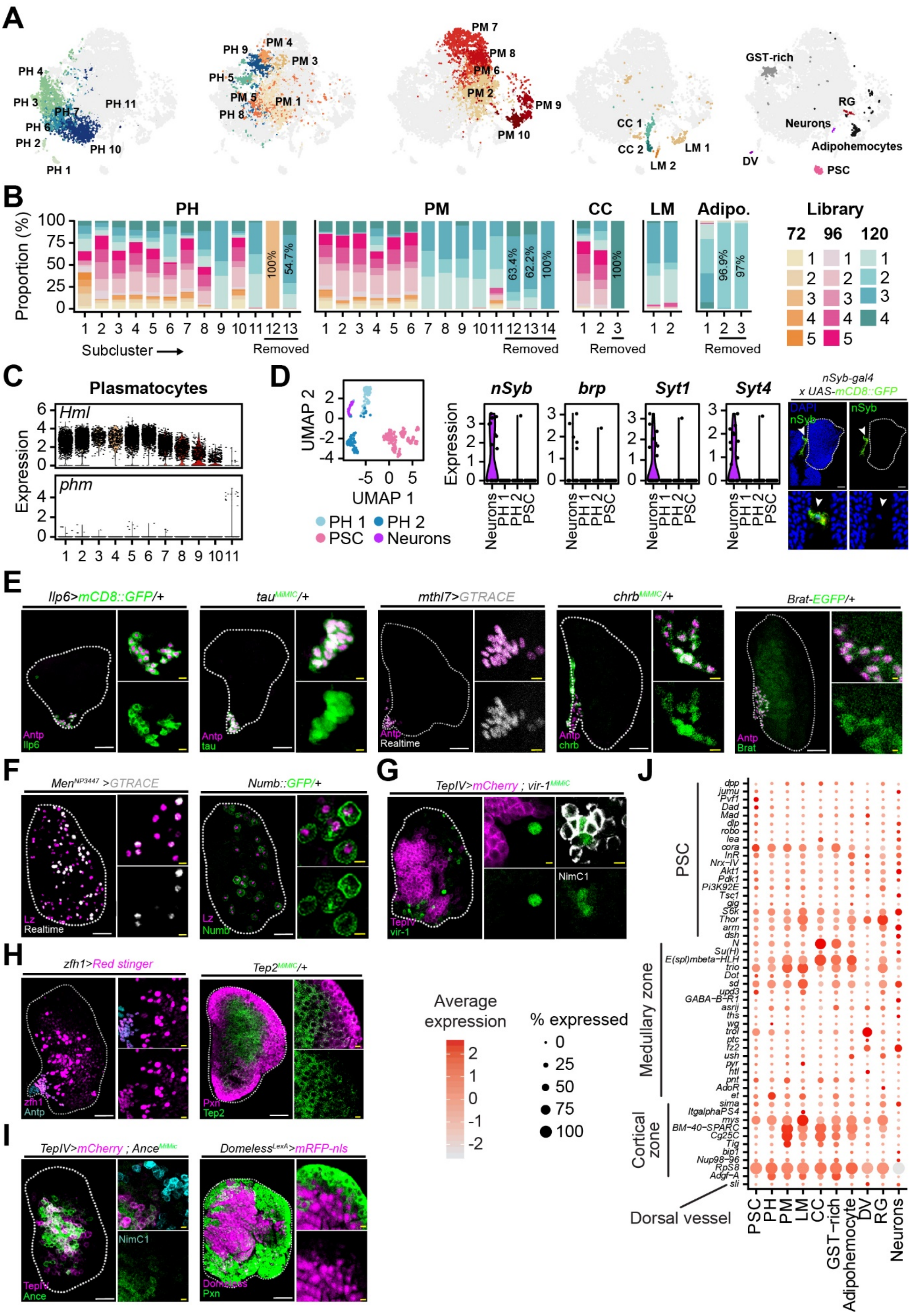
A
PH 1
PH 2
PH 3
PH 4
PH 6
PH 7
PH 10
PH 11
PM 1
PM 3
PM 4
PM 5
PH 5
PH 8
PH 9
PM 2
PM 6
PM 7
PM 8
PM 9
PM 10
CC 1
CC 2
LM 1
LM 2
GST-rich
RG
Neurons
Adipohemocytes
DV
PSC
B
PH
PM
CC
LM
Adipo.
Library
72
96
120
Proportion (%)
Subcluster
Removed
100%
54.7%
63.4%
62.2%
96.9%
97%
C
Plasmatocytes
Hml
phm
Expression
D
UMAP 2
UMAP 1
PH 1
PH 2
PSC
Neurons
nSyb
brp
Syt1
Syt4
nSyb-gal4
x UAS-mCD8::GFP
DAPI
E
Ilp6>mCD8::GFP/+
tauMiMIC/+
mthl7>GTRACE
chrbMiMIC/+
Brat-EGFP/+
Antp
Ilp6
tau
Realtime
chrb
Brat
F
MenNP3447 >GTRACE
Numb::GFP/+
Lz
Numb
G
TepIV>mCherry ; vir-1MiMIC
NimC1
TepIV
vir-1
H
zfh1>Red stinger
Tep2MiMIC/+
zfh1
Pxn
Tep2
I
TepIV>mCherry ; AnceMiMIC
DomelessLexA>mRFP-nls
Ance
Domeless
J
Average expression
% expressed
PSC
Medullary zone
Cortical zone
Dorsal vessel
dpp
jumu
Pvf1
Dad
Mad
dlp
robo
lea
cora
InR
Nrx-IV
Akt1
Pdk1
Pi3K92E
Tsc1
gig
S6k
Thor
arm
dsh
N
Su(H)
E(spl)mbeta-HLH
trio
Dot
sd
upd3
GABA-B-R1
asrij
ths
wg
trol
ptc
fz2
ush
pyr
htl
pnt
AdoR
et
sima
ItgalphaPS4
mys
BM-40-SPARC
Cg25C
Tig
bip1
Nup98-96
RpS8
Adgf-A
sli
GST-rich
Adipohemocyte
DV
RG
Neurons

**Figure S2. Filtration of library-specific subclusters and *in vivo* validation of novel or known marker genes in the lymph gland**

(A) Subclusters exclusive to prohemocytes (PHs), intersection of prohemocytes and plasmatocytes (PHs and PMs), exclusive plasmatocytes (PMs), crystal cells or lamellocytes (CCs or LMs), other hemocytes (GST-rich, adipohemocytes, and the PSC), and non-hematopoietic cells (neurons, dorsal vessel (DV), and ring gland (RG), in *t*-SNE plots.

(B) Proportion of sequencing libraries per one subcluster found in major cell types. Underlined subclusters are filtered out in the subsequent analyses.

(C) Expression of *phm*, a ring gland marker, in the PM11 subcluster. *Hml* is barely expressed in PM11, and accordingly renamed as ring gland subcluster.

(D) Identification of neuronal cells from subclustering analyses of PH1, PH2, and PSC (left). Expression of neuronal marker genes—*nSyb*, *brp*, *Syt1*, and *Syt4*— in the prospective neuron subcluster (middle). The presence of neuronal projections near the lymph gland (green, *nSyb*; blue, DAPI; *nSyb-Gal4 UAS-mCD8::GFP*; right).

(E) Novel PSC markers in the lymph gland. Antp$^+$ PSC cells (magenta) co-localize with *Ilp6* (green; *Ilp6-Gal4 UAS-mCD8::GFP*), *tau* (green; *tau*$^{MiMiC}$), *mthl7* (white; *mthl7-Gal4 UAS-GTRACE*), *chrb* (green; *chrb*$^{MiMiC}$), or *Brat* (green; *Brat-GFP*). Magnified images show co-localization of Antp and the new markers (right in each panel).

(F) Novel crystal cell markers in the lymph gland. Lz$^+$ crystal cells (magenta) co-localized with *Men* (white; *Men-Gal4 UAS-GTRACE*) or *Numb* (green; *Numb::GFP*). Lz co-localizes with a subset of *Men* while all the *Numb*$^+$ and Lz$^+$ overlap. Magnified images show co-localization of Lz and the new markers (right in each panel).

(G) The novel plasmatocyte marker, *vir-1* (green; *vir-1*$^{MiMiC}$) does not co-localize with *TepIV*$^{+}$ prohemocytes (magenta; left), but overlaps with NimC1 (white; left). Magnified images show co-localization of *vir-1* with NimC1 (right in each panel).

(H) Newly identified prohemocyte markers. *zfh1* (magenta; *zfh1-Gal4 UAS-Red singer*; left) co-localizes in both Antp$^{+}$ PSC (cyan) and prohemocytes. *Tep2* (green; *Tep2*$^{MiMiC}$; right) partially co-localizes with Pxn (magenta). Magnified images in each panel show co-localization of *zfh1* (magenta; left) and Antp (cyan; left) or *Tep2* (green, left) and Pxn (magenta, right) (right in each panel).

(I) Newly identified or generated prohemocyte markers. *TepIV*$^{+}$ prohemocytes (magenta; left) co-localizes with *Ance* (green; *TepIV-Gal4 UAS-mCherry Ance*$^{MiMIC}$) but is separable from NimC1$^{+}$ mature PMs (cyan). *Domeless-LexA* generated in this study (magenta; *Domeless-LexA LexAop-mRFP-nls*) partially co-localizes with Pxn (green). Magnified images show co-localization of *Ance* (green), *TepIV* (magenta), and NimC1 (cyan) or *Domeless* (magenta) and Pxn (green) (right in each panel).

(J) Dot plot showing the levels of marker genes specific to the PSC, the medullary zone, the cortical zone, or the dorsal vessel. Additional genes are indicated in Figure 2B.

White scale bar is 30 μm. Lymph gland is demarcated by white dotted line. Arrowhead indicates a neuron. Magnified images are shown at the bottom (right).

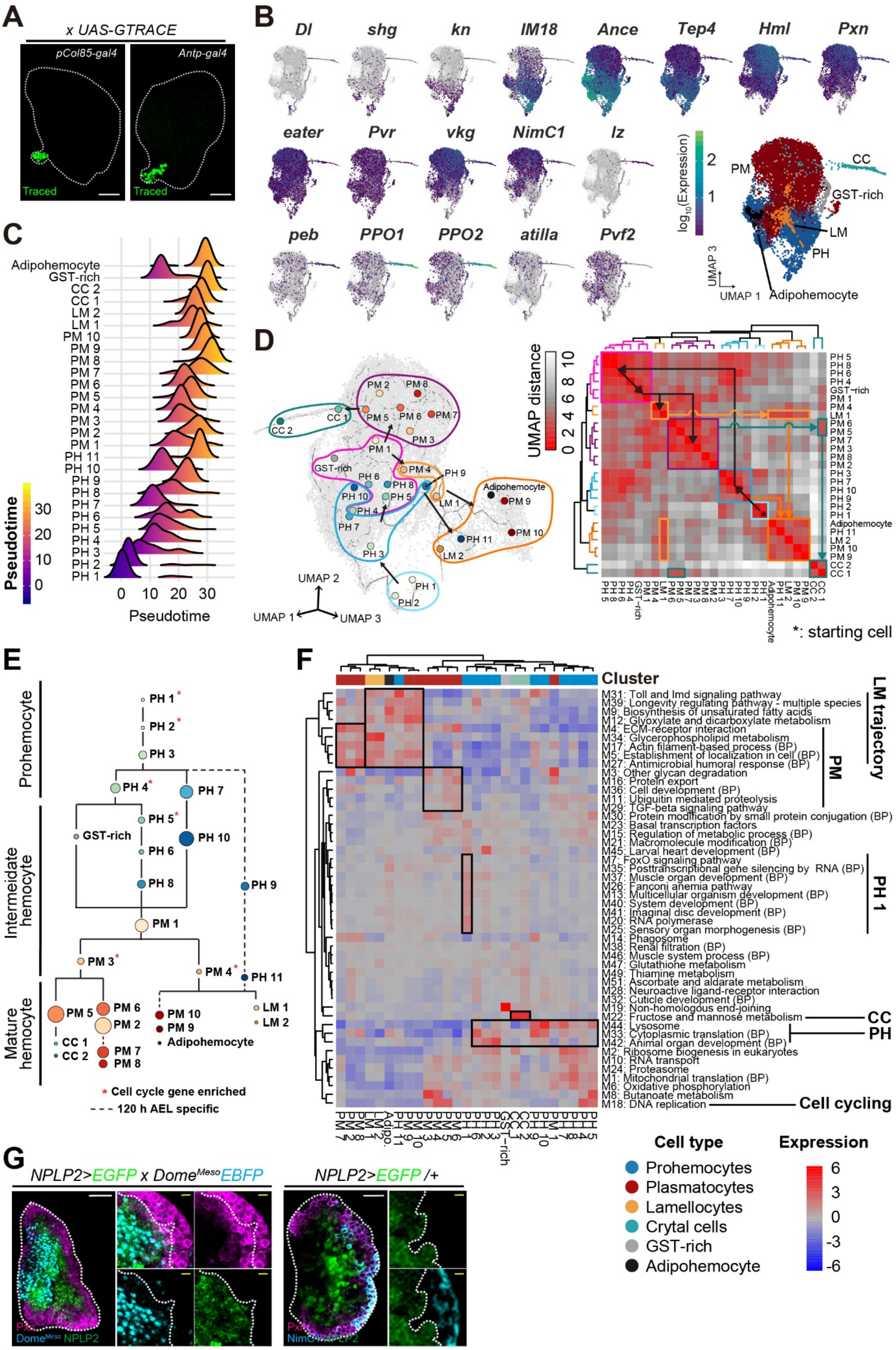
A
x UAS-GTRACE
pCol85-gal4
Antp-gal4
Traced
Traced
B
Dl
shg
kn
IM18
Ance
Tep4
Hml
Pxn
eater
Pvr
vkg
NimC1
lz
peb
PPO1
PPO2
atilla
Pvf2
log10(Expression)
2
1
PM
CC
GST-rich
LM
PH
Adipohemocyte
UMAP 3
UMAP 1
C
Adipohemocyte
GST-rich
CC 2
CC 1
LM 2
LM 1
PM 10
PM 9
PM 8
PM 7
PM 6
PM 5
PM 4
PM 3
PM 2
PM 1
PH 11
PH 10
PH 9
PH 8
PH 7
PH 6
PH 5
PH 4
PH 3
PH 2
PH 1
Pseudotime
30
20
10
0
0 10 20 30
Pseudotime
D
UMAP 2
UMAP 1
UMAP 3
UMAP distance
0 2 4 6 8 10
*: starting cell
E
Prohemocyte
Intermediate hemocyte
Mature hemocyte
PH 1*
PH 2*
PH 3
PH 4*
PH 7
PH 5*
GST-rich
PH 10
PH 6
PH 8
PH 9
PM 1
PM 3*
PM 4*
PH 11
PM 5
PM 6
PM 2
CC 1
CC 2
PM 7
PM 8
PM 10
PM 9
Adipohemocyte
LM 1
LM 2
* Cell cycle gene enriched
- - - 120 h AEL specific
F
Cluster
M31: Toll and Imd signaling pathway
M39: Longevity regulating pathway - multiple species
M9: Biosynthesis of unsaturated fatty acids
M12: Glyoxylate and dicarboxylate metabolism
M4: ECM-receptor interaction
M34: Glycerophospholipid metabolism
M17: Actin filament-based process (BP)
M5: Establishment of localization in cell (BP)
M27: Antimicrobial humoral response (BP)
M3: Other glycan degradation
M16: Protein export
M36: Cell development (BP)
M11: Ubiquitin mediated proteolysis
M29: TGF-beta signaling pathway
M30: Protein modification by small protein conjugation (BP)
M23: Basal transcription factors
M15: Regulation of metabolic process (BP)
M21: Macromolecule modification (BP)
M45: Larval heart development (BP)
M7: FoxO signaling pathway
M35: Posttranscriptional gene silencing by RNA (BP)
M37: Muscle organ development (BP)
M26: Fanconi anemia pathway
M13: Multicellular organism development (BP)
M40: System development (BP)
M41: Imaginal disc development (BP)
M20: RNA polymerase
M25: Sensory organ morphogenesis (BP)
M14: Phagosome
M38: Renal filtration (BP)
M46: Muscle system process (BP)
M47: Glutathione metabolism
M49: Thiamine metabolism
M51: Ascorbate and aldarate metabolism
M28: Neuroactive ligand-receptor interaction
M32: Cuticle development (BP)
M19: Non-homologous end-joining
M22: Fructose and mannose metabolism
M44: Lysosome
M33: Cytoplasmic translation (BP)
M42: Animal organ development (BP)
M2: Ribosome biogenesis in eukaryotes
M10: RNA transport
M24: Proteasome
M1: Mitochondrial translation (BP)
M6: Oxidative phosphorylation
M8: Butanoate metabolism
M18: DNA replication
LM trajectory
PM
PH 1
CC
PH
Cell cycling
Cell type
Prohemocytes
Plasmatocytes
Lamellocytes
Crytal cells
GST-rich
Adipohemocyte
Expression
6
3
0
-3
-6
G
NPLP2>EGFP x Dome^Meso EBFP
NPLP2>EGFP /+
Pxn
Dome^Meso NPLP2

**Figure S3. Molecular features and distribution of subclusters along developmental pseudotime trajectories**

(A) PSC cells do not contribute to the rest of the lymph gland. Either *pCol85-Gal4* or *Antp-Gal4* is not traced beyond the PSC (green, traced; *pCol85-Gal4 UAS-GTRACE* or *Antp-Gal4 UAS-GTRACE*). Scale bar, 30 μm. Lymph glands are demarcated by white dotted lines.

(B) Expression patterns of 18 known marker genes—*Dl, shg, kn (col), IM18, Ance, Tep4, Hml, Pxn, eater, Pvr, vkg, NimC1, lz, peb, PPO1, PPO2, atilla, and Pvf2*—of major cell types in two-dimensional (UMAP 1 and 3) pseudotime trajectories. Distribution of cell types is shown at the bottom-right. Color bar denotes the $\log_{10}$-scaled level of gene expression. Grey color indicates no expression.

(C) Relative densities of subclusters along pseudotime. PH1 and PH2 emerge at the earliest pseudotime. PH3-PH8 and PH10 span through the mid-pseudotime while PH9 and PH11 peak at later time point. PM9-PM10, LM1, LM2, CC2, and adipohemocytes are the last cell types to differentiate.

(D) Distribution of subclusters in three-dimensional pseudotime trajectory (left), and clustering of distances between subclusters (right). Subclusters are grouped according to distances and directions of differentiation (left). PH1 and PH2 (white box) are the earliest subclusters among prohemocytes. PH11, PM9, PM10, adipohemocytes, LM1, and LM2 (orange box) form a separate group. PM5 is the closest to CC1 and CC2 (green box), whereas PM4 shares similarities with the lamellocyte lineages (orange box). Color bar denotes the UMAP distance (right).

(E) Simplified stick-and-ball presentation of the trajectory. Red asterisks indicate subclusters with high expressions of cell cycle genes. Dotted lines show lineages that appear only at 120 h AEL. Size of the ball represents subcluster proportion. Each subcluster color is identical to Supplementary Figure 3D.

(F) Heatmap showing the level of gene expression modules of subclusters. Gene modules clustered in a specific cell type are indicated in black boxes. The column annotation indicates major cell types used in the analysis.

(G) *Nplp2* (green) partially co-localizes with *Dome*$^{Meso+}$ prohemocytes (cyan) or with Pxn$^{+}$ plasmatocytes (magenta) (left; *Nplp2-Gal4 UAS-EGFP; Dome*$^{Meso}$*-EBFP*). However, *Nplp2* (green) does not overlap with a mature plasmatocyte marker, NimC1 (cyan) (right; *Nplp2-Gal4 UAS-EGFP*). Magnified images are shown at the right of each panel. Yellow dotted line demarcates the margins of *Nplp2*. White scale bar, 30 μm. Yellow scale bar, 3 μm. Lymph glands are demarcated by white dotted lines.

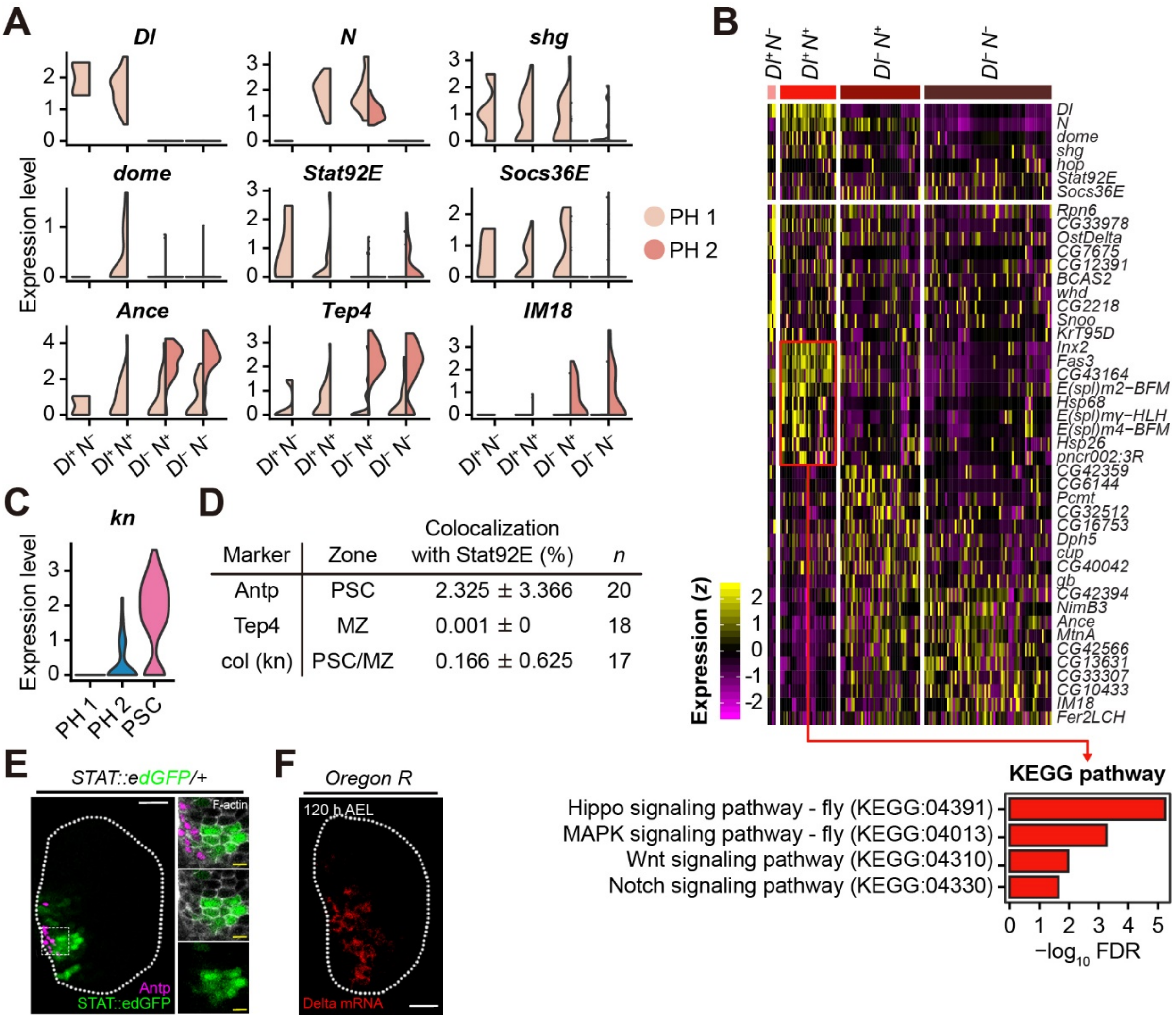


**Figure S4. PH1 cells express Dl and JAK/STAT**

(A) Expression of PH1- or prohemocyte marker genes in $Dl^{+}N^{-}$, $Dl^{+}N^{+}$, $Dl^{-}N^{+}$, or $Dl^{-}N^{-}$ PH1. Expression values of PH1 (light brown) and PH2 (brown) are shown separately.

(B) Marker genes classified by the differential expression of *Dl* and *N* in PH1 and PH2. $Dl^{+}N^{-}$ cells exhibit high levels of *Dl*, *shg*, *STAT92E*, and *Socs36E*. $Dl^{+}N^{+}$ cells show gradual decrease or increase in *Dl* or *N* and express *ena*, *bowl*, *DnaJ-1*, *EcR*, *br*, and *Argk*. $Dl^{-}N^{+}$ cells display *MED27*, *Atg8a*, and *RhoGDI* along with high *N*. $Dl^{-}N^{-}$ cells are distinguishable from other cell types and express high *Ance*. $Dl^{+}N^{+}$ cells show enriched Hippo, MAPK,

Wnt, and Notch signaling pathways based on KEGG pathway analysis. Color bar indicates the level of scaled gene expression.

(C) Violin plots indicating differential levels of *kn* (*col*) in PH1, PH2 and in the PSC.

(D) Quantitation of co-localization of *Stat92E::edGFP* and representative marker genes, *Antp*, *Tep4*, or *col*. *Antp*$^+$ PSC, *Tep4*$^+$ MZ or *col*$^+$ PSC/MZ cells do not show significant overlap with *Stat92E::edGFP*. *n* indicates the number of samples.

(E) *STAT92E*$^+$ PH1 cells (green) physically interact with Antp$^+$ (magenta) PSC cells (left). When visualized with F-actin (white), Antp$^+$ cells and *STAT92E*$^+$ cells share F-actin membrane but do not co-localize (right). White scale bar, 30 μm. Yellow scale bar, 3 μm. Lymph glands are demarcated by white dotted lines.

(F) Fluorescent *in situ* hybridization against *Dl* mRNA (red) in the lymph gland at 120 h AEL. *Dl* is localized in the medioposterior region of the lymph gland. Scale bar, 30 μm. Lymph gland is demarcated by white dotted line.

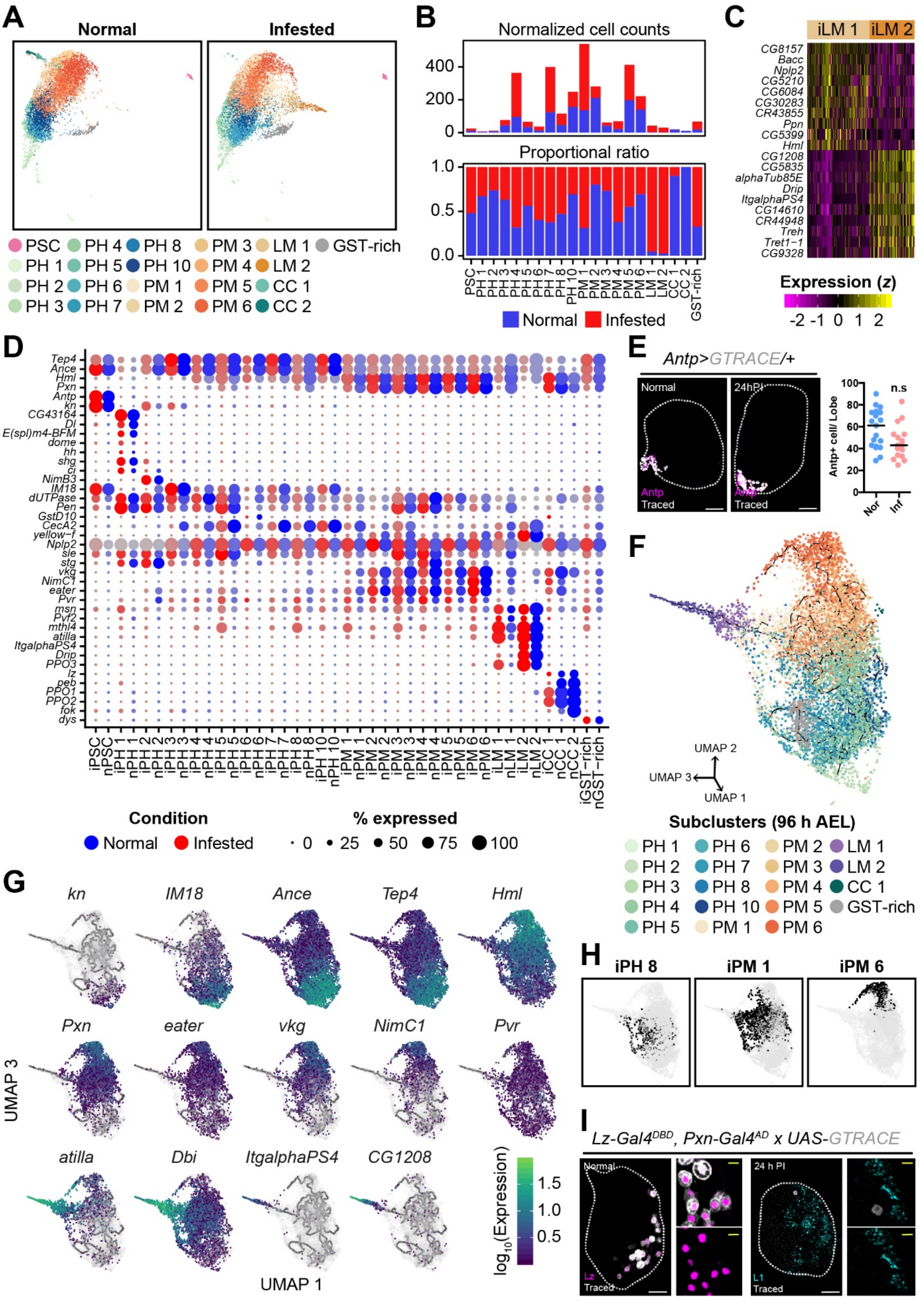

A
Normal
Infested
PSC
PH 1
PH 2
PH 3
PH 4
PH 5
PH 6
PH 7
PH 8
PH 10
PM 1
PM 2
PM 3
PM 4
PM 5
PM 6
LM 1
LM 2
CC 1
CC 2
GST-rich
B
Normalized cell counts
Proportional ratio
Normal
Infested
C
iLM 1
iLM 2
CG8157
Bacc
Nplp2
CG5210
CG6084
CG30283
CR43855
Ppn
CG5399
Hml
CG1208
CG5835
alphaTub85E
Drip
ItgalphaPS4
CG14610
CR44948
Treh
Tret1−1
CG9328
Expression (z)
D
Tep4
Ance
Hml
Pxn
Antp
kn
CG43164
Dl
E(spl)m4-BFM
dome
hh
shg
ci
NimB3
IM18
dUTPase
Pen
GstD10
CecA2
yellow−f
Nplp2
sle
stg
vkg
NimC1
eater
Pvr
msn
Pvf2
mthl4
atilla
ItgalphaPS4
Drip
PPO3
lz
peb
PPO1
PPO2
fok
dys
Condition
Normal
Infested
% expressed
E
Antp>GTRACE/+
Normal
24hPI
Antp
Traced
Antp+ cell/ Lobe
n.s
Nor
Inf
F
UMAP 2
UMAP 3
UMAP 1
Subclusters (96 h AEL)
G
kn
IM18
Ance
Tep4
Hml
Pxn
eater
vkg
NimC1
Pvr
atilla
Dbi
ItgalphaPS4
CG1208
UMAP 3
UMAP 1
log10(Expression)
H
iPH 8
iPM 1
iPM 6
I
Lz-Gal4DBD, Pxn-Gal4AD x UAS-GTRACE
Normal
Lz
Traced
24 h PI
L1
Traced

**Figure S5. Comparison of normal and wasp infested lymph gland datasets and pseudotime trajectory analysis of wasp infested lymph glands**

(A) Two-dimensional projections of subclustered cells in normal (left) and wasp infested (right) lymph glands. Upon wasp infestation, late prohemocyte, early plasmatocyte, GST-rich, and lamellocyte cell population expand while crystal cells are significantly reduced. The PSC remains intact. Colors indicate each subcluster.

(B) Normalized cell counts (top) or proportional ratio (bottom) of subclustered cells in normal (blue) or wasp infested (red) lymph gland hemocytes.

(C) 20 top genes in iLM1 or iLM2. iLM1 expresses the *Nplp2* and *Hml* marker genes for intermediate cells and plasmatocytes, respectively. iLM2 expresses *alphaTub85E*, *Drip*, and *ItgalphaPS4*. Color bar indicates the level of scaled gene expression.

(D) Dot plot expressing the level of known and novel marker genes in normal (blue) or wasp infested (red) lymph glands (prefix 'n' and 'i' for normal and wasp infestation, respectively).

(E) PSC cell numbers do not change upon wasp infestation (Antp, magenta; traced, white; *Antp-Gal4 UAS-GTRACE*). Graph indicates the number of Antp$^+$ cells per one lymph gland lobe. Scale bar, 30 μm. Lymph gland is demarcated by white dotted line.

(F) Three-dimensional pseudotime trajectory of subclustered cells from the normal lymph gland dataset.

(G) Expression patterns of 14 known marker genes– *kn* (*col*), *IM18*, *Ance*, *Tep4*, *Hml*, *Pxn*, *eater*, *vkg*, *NimC1*, *Pvr*, *atilla*, *Dbi*, *ItgalphaPS4*, and *CG1208*–in two-dimensional (UMAP 1 and 3) pseudotime trajectory. Color bar denotes the $\log_{10}$-scaled level of gene expression. Grey color indicates no expression.

(H) Three subclusters directly associate with the lamellocyte lineages under wasp infestation. iPH8 and iPM1 are the largest subclusters that give rise to iLMs. These are implicated as intermediate populations in Supplementary Figure 2A and 3D. Another origin of iLM is iPM6, the most mature PMs at 96 h AEL.

(I) iLM and iCC lineages are separable. When differentiating crystal cells are traced after wasp infestation, iLMs emerge independently of iCCs (*Lz-Gal4$^{DBD}$, Pxn-Gal4$^{AD}$ UAS-GTRACE*). *Lz-Gal4$^{DBD}$, Pxn-Gal4$^{AD}$ UAS-GTRACE* trace the majority of *Lz*-expressing crystal cells (Lz, magenta; traced, white). However, L1$^{+}$ iLMs are *Lz$^{DBD}$/Pxn$^{AD}$*-negative (L1, cyan; traced, white). White scale bar, 30 μm. Yellow scale bar, 3 μm. Lymph glands are demarcated by white dotted lines. Magnified images are shown on the right of each panel.

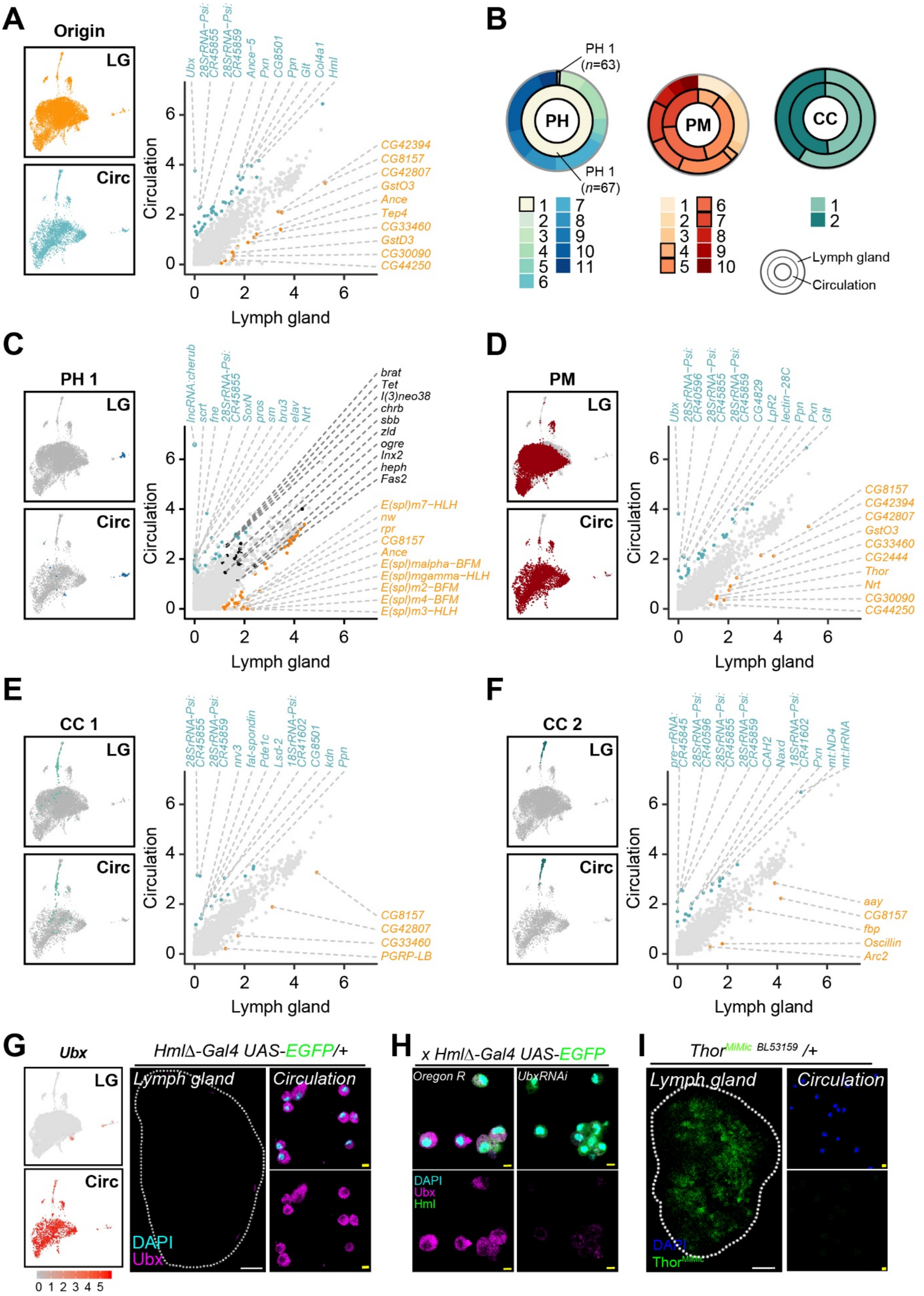

A
Origin
LG
Circ
Ubx
28SrRNA-Psi:CR45855
28SrRNA-Psi:CR45859
Ance-5
Pxn
CG8501
Ppn
Glt
Col4a1
Hml
CG42394
CG8157
CG42807
GstO3
Ance
Tep4
CG33460
GstD3
CG30090
CG44250
Circulation
Lymph gland
B
PH 1 (n=63)
PH 1 (n=67)
PH
PM
CC
Lymph gland
Circulation
C
PH 1
lncRNA:cherub
scrt
fne
28SrRNA-Psi:CR45855
SoxN
pros
sm
bru3
elav
Nrt
brat
Tet
l(3)neo38
chrb
sbb
zld
ogre
Inx2
heph
Fas2
E(spl)m7-HLH
nw
rpr
CG8157
Ance
E(spl)malpha-BFM
E(spl)mgamma-HLH
E(spl)m2-BFM
E(spl)m4-BFM
E(spl)m3-HLH
D
PM
Ubx
28SrRNA-Psi:CR40596
28SrRNA-Psi:CR45855
28SrRNA-Psi:CR45859
CG4829
LpR2
lectin-28C
Ppn
Pxn
Glt
CG8157
CG42394
CG42807
GstO3
CG33460
CG2444
Thor
Nrt
CG30090
CG44250
E
CC 1
28SrRNA-Psi:CR45855
28SrRNA-Psi:CR45859
nrv3
fat-spondin
Pde1c
Lsd-2
18SrRNA-Psi:CR41602
CG8501
kdn
Ppn
CG8157
CG42807
CG33460
PGRP-LB
F
CC 2
pre-rRNA:CR45845
28SrRNA-Psi:CR40596
28SrRNA-Psi:CR45855
28SrRNA-Psi:CR45859
CAH2
Naxd
18SrRNA-Psi:CR41602
Pxn
mt:ND4
mt:lrRNA
aay
CG8157
fbp
Oscillin
Arc2
G
Ubx
HmlΔ-Gal4 UAS-EGFP/+
Lymph gland
Circulation
DAPI
Ubx
H
x HmlΔ-Gal4 UAS-EGFP
Oregon R
UbxRNAi
DAPI
Ubx
Hml
I
Thor MiMic BL53159 /+
Lymph gland
Circulation
DAPI
Thor MiMic

**Figure S6. Lineage-specific cell types and marker genes for the lymph gland- or the embryonically derived hemocytes**

(A) Comparison of gene expressions between the lymph gland and circulating hemocytes. Top ranked lymph gland- (right) or circulation-specific (top) differentially expressed genes are indicated in colored gene names (orange, upregulated in lymph glands; cyan, upregulated in circulation).

(B) Pie charts showing the relative proportion of subclusters in the lymph gland and in circulation datasets. The outer graph indicates proportions of the lymph gland and the inner graph shows those of the circulation dataset. Demarcated subclusters indicate those found in circulation.

(C-F) Comparison of gene expressions between the lymph gland and in circulation. PH1 (C), PM (D), CC1 (E), and CC2 (F). Top ranked lymph gland- (right) or circulation-specific (top) differentially expressed genes are indicated in colored gene names (orange, upregulated in lymph glands; cyan, upregulated in circulation).

(G) Expression of *Ubx* is exclusive in circulating plasmatocytes. Comparison of *Ubx* levels between the lymph gland and in circulation (left). The lymph gland does not show *Ubx* expression (cyan, DAPI; left). Circulating plasmatocytes display *Ubx* (cyan, DAPI; magenta, Ubx; right) (*Hml-Gal4 UAS-EGFP* control). White scale bar indicates 30 μm. White dotted line demarcates the lymph gland.

(H) Verification of *Ubx* expression in circulating hemocytes. Expression levels of *Ubx* in circulating plasmatocytes are significantly reduced when RNAi against *Ubx* is expressed (green, Hml; cyan, DAPI; magenta, Ubx) (*Hml-Gal4 UAS-EGFP UAS-UbxRNAi*). Yellow scale bar indicates 3 μm.

(I) Confirmation of *Thor* expression in the lymph gland. Comparison of *Thor* levels between the lymph gland (left) and in circulation (right). The lymph gland expresses *Thor* (green, Thor; left) (*Thor*$^{MiMiC}$/+) while circulating plasmatocytes do not express *Thor* (blue, DAPI; green, Thor; right). White scale bar indicates 30 μm, yellow scale bar indicates 3 μm. White dotted line demarcates the lymph gland.

**Supplementary Table S1. Basic statistics of scRNA-seq**

(A) Basic statistics of lymph grand scRNA-seq libraries. (B) Basic statistics of scRNA-seq libraries from wasp-infested lymph glands. (C) Basic statistics of scRNA-seq libraries from circulation hemocytes.

**Supplementary Table S2. Signature genes for cell types and subclusters**

(A) Signature genes for ten hemocytes and non-hematopoietic cell types. (B) Signature genes for 31 hematocytes and non-hematopoietic subclusters.

**Supplementary table S3. Putative *Drosophila* markers screened in this study**

(A) Putative *Drosophila* markers screened in this study. (B) Putative markers for embryonic and lymph gland hemocytes. All Gal4 lines are crossed with *UAS-GTRACE*. MiMiC lines are confirmed with GFP antibody.

**Supplementary table S4. *Drosophila* and Human orthologs**

(A) Top 30 conserved signature genes. (B) Top conserved signature genes for wasp infestation data.

**Supplementary video 1. Three-dimensional trajectory landscape**
